# *Toxoplasma gondii* F-Box Protein L2 Silences Feline-Restricted Genes Necessary for Sexual Commitment

**DOI:** 10.1101/2023.12.18.572150

**Authors:** Carlos G. Baptista, Sarah Hosking, Elisabet Gas-Pascual, Loic Ciampossine, Steven Abel, Mohamed-Ali Hakimi, Victoria Jeffers, Karine Le Roch, Christopher M. West, Ira J. Blader

## Abstract

*Toxoplasma gondii* is a foodborne pathogen that can cause severe and life-threatening infections in fetuses and immunocompromised patients. Felids are its only definitive hosts, and a wide range of animals, including humans, serve as intermediate hosts. When the transmissible bradyzoite stage is orally ingested by felids, they transform into merozoites that expand asexually, ultimately generating millions of gametes for the parasite sexual cycle. However, bradyzoites in intermediate hosts differentiate exclusively to disease-causing tachyzoites, which rapidly disseminate throughout the host. Though tachyzoites are well-studied, the molecular mechanisms governing transitioning between developmental stages are poorly understood. Each parasite stage can be distinguished by a characteristic transcriptional signature, with one signature being repressed during the other stages. Switching between stages requires substantial changes in the proteome, which is achieved in part by ubiquitination. F-box proteins mediate protein poly-ubiquitination by recruiting substrates to SKP1, Cullin-1, F-Box protein E3 ubiquitin ligase (SCF-E3) complexes. We have identified an F-box protein named *Toxoplasma gondii* F-Box Protein L2 (TgFBXL2), which localizes to distinct nuclear sites. TgFBXL2 is stably engaged in an SCF-E3 complex that is surprisingly also associated with a COP9 signalosome complex that negatively regulates SCF-E3 function. At the cellular level, TgFBXL2-depleted parasites are severely defective in centrosome replication and daughter cell development. Most remarkable, RNA seq data show that TgFBXL2 conditional depletion induces the expression of genes necessary for sexual commitment. We suggest that TgFBXL2 is a latent guardian of sexual stage development in *Toxoplasma* and poised to remove conflicting proteins in response to an unknown trigger of sexual development.

**AUTHOR SUMMARY:** *Toxoplasma gondii* is a protozoan parasite that replicates sexually in felids and asexually in nearly all other mammals with each life stage having a specific transcriptional profile. When life stage specific transcription is not properly controlled, the parasite dies and therefore it’s important to understand what inhibits expression of sexual stage genes during asexual growth and vice versa. Here we identify a ubiquitin E3 ligase complex that inhibits sexual stage gene expression during asexual growth.

## INTRODUCTION

*Toxoplasma gondii* is an intracellular apicomplexan parasite that is responsible for one of the world’s most widespread parasitic infections; an estimated one-third of humans worldwide are infected [1]. *Toxoplasma* infections can occur by ingesting either of the parasite’s transmissible forms: bradyzoites that reside inside tissue cysts or sporozoite-laden oocysts [2, 3]. Upon ingestion by an intermediate host, the parasite differentiates into tachyzoites, the rapidly dividing and highly motile form of *Toxoplasma*, which disseminate rapidly throughout the host’s body [4, 5]. In response to the host’s immune system, the parasite forms bradyzoite-containing tissue cysts, evading elimination and establishing chronic infection [4, 6]. However, for individuals who are unable to mount an appropriate immune response, *Toxoplasma* can cause debilitating and life-threatening disease [1, 7] via uncontrolled parasite growth and immune-mediated tissue damage [6, 8–13].

Although the parasite has a remarkable ability to infect a diverse range of warm-blooded animals as intermediate hosts, felids are the only definitive host [14]. However, when domestic cats and other felids ingest tissue cysts the parasite differentiates into merozoites that replicates by a very distinct process named merogony [15]. Several rounds of asexual division and amplification are followed by differentiation into macro- and microgamonts [16, 17]. Fertilization results in immature oocysts that are shed with the feces into the environment where they become infectious over the course of several days [18, 19].

Tachyzoites and merozoites can be distinguished by their respective transcriptional signatures, with each being repressed during the other stage [20]. Recent advances in understanding *Toxoplasma* genetic reprogramming have uncovered several transcriptional factors and epigenetic modifiers, yet the mechanisms linking gene expression and stage transitions remain poorly understood [21–23]. Switching between stages requires significant changes in protein profiles mediated in part by posttranslational modifications such as poly-ubiquitination, which mediates a substantial fraction of protein turnover and cell differentiation [24, 25].

F-box proteins (FBPs) are critical elements in several processes, including cellular differentiation, transcription, and cell cycle progression [26–29]. FBPs are a family of proteins defined by the presence of an F-box domain, a region of about 40 amino acids that docks with SKP1 [30, 31]. FBPs mediate protein ubiquitination by recruiting substrates to SKP1/Cullin-1/FBP/RBX1 containing E3 ubiquitin ligase (SCF-E3) complexes [32–38]. Previously, we identified 18 putative FBPs in the *Toxoplasma* genome and a CRISPR screen indicated that the *Toxoplasma* F-box Protein L2 (TgFBXL2) is the FBP most required for parasite fitness [39, 40]. TgFBXL2 is one of two L-type F-box proteins in *Toxoplasma* that have a C-terminal region comprising a series of leucine-rich repeats (LRRs). LRRs are commonly found in FBPs of other organisms where they mediate contact with substrates for poly-ubiquitination.

Here, we report that TgFBXL2 localizes to a distinct nuclear site and is essential for tachyzoite growth due to its role in regulating centrosome replication and daughter cell biogenesis. Most remarkably, TgFBXL2 conditional depletion induces the expression of genes involved in sexual commitment such as merozoite restricted surface-related genes (SRGs) and GRA80. Interestingly, our RNA seq data do not indicate down-regulation of tachyzoite-expressed genes, indicating TgFBXL2 conditional depletion mostly promotes activation of pre-sexual and sexual transcription in tachyzoites.

## RESULTS

### Identification of TgFBXL2 as a SCF-E3 Subunit

Earlier, we identified candidate F-box proteins by interrogating the *Toxoplasma* genome and TgSKP1 interactome and identified TgFBXL2 with medium confidence as a candidate TgSKP1 interacting protein [40]. TgFBXL2 (TGGT1_313200) contains 832 amino acids and has 12 LRR sequence motifs that are predicted to fold into a solenoid-like structure characteristic of substrate receptor domains found in many FBPs **(Fig. S1)**. Upstream of the LRR motifs lies an F-box-like sequence that likely explains its presence in the TgSKP1 interactome [40]. Upstream of the F-box motif are two predicted nuclear localization signals. These sequence motifs are conserved in most Apicomplexa; however, the F-box motif sequence and the NLS motifs diverge dramatically in non-Sarcocystid species, including *Plasmodium spp*. The remainder of the TgFBXL2 sequence is very poorly conserved even in *Neospora caninum*.

Because TgFBXL2 was found using a genome-wide CRISPR screen to be important for parasite fitness, we created a TgFBXL2 conditional expression strain by replacing the TgFBXL2 endogenous promoter in the TATiΔKu80 strain with an anhydrotetracycline (ATC)-responsive promoter and an amino-terminal 3xHA epitope tag cloned in frame to generate the ^HA^TgFBXL2 strain [41] (**Fig. 1A&B**). Western blotting lysates from ^HA^TgFBXL2-expressing parasites but not the parental TATiΔKu80 strain with anti-HA antibodies revealed a single immunoreactive band at ~100 kDa, which is the approximate expected molecular weight of ^HA^TgFBXL2 **(Fig. 1C; Input)**. Immunoprecipitating ^HA^TgFBXL2 with anti-HA and Western blot detection of TgSKP1 in the immunoprecipitates confirmed TgFBXL2/TgSKP1 interactions **(Fig. 1C; IP)**.

**Figure 1.**
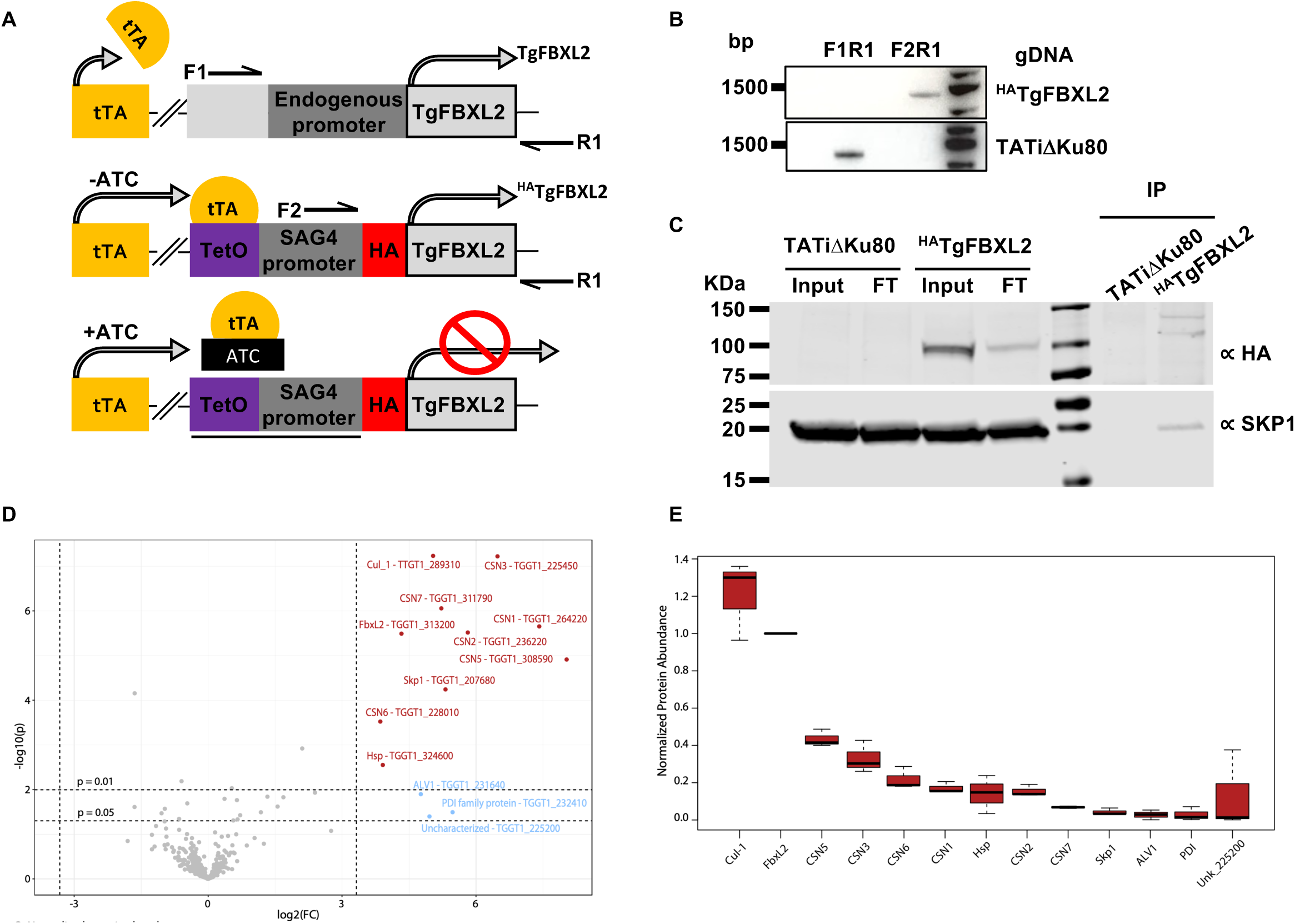
TgFBXL2 is a *Toxoplasma* F-Box Protein. **(A).** Schematic illustration of ^HA^TgFBXL2 anhydrotetracycline (ATC)-mediated gene expression. The endogenous promoter of TgFBXL2 was replaced with a SAG4 promoter construct in which a tetracycline transactivator (tTA) binding element was cloned upstream. Addition of ATC reduces transcription by preventing tTA binding to the tetracycline responsive promoter. **(B).** PCR of ^HA^TgFBXL2 genomic DNA showing correct integration of anhydrotetracycline responsive promoter using primers F2 and R1 as show in A. **(C).** Lysates prepared from ^HA^TgFBXL2 and parental TATiΔKu80 were either Western blotted (Input) or incubated with anti-HA antibodies (IP) that were captured by Protein G Sepharose. ^HA^TgFBXL2 and TgSKP1 were detected in lysates (input), flow through (FT), and immunoprecipitates (IP) by Western blotting with anti-HA and TgSKP1 antibodies. **(D).** Volcano plot of identified proteins in large-scale lysates immunoprecipitated with anti-HA beads and analyzed by MS. Proteins in red were 10-fold enriched from the ^HA^TgFBXL2 strain vs. parental with *P*-value ≤0.01 and therefore are considered *bona-fide* TgFBXL2 interactors. Relaxing the p-value to ≤0.05 resulted in 3 additional proteins that are likely to be false positives. See Table S1 for complete dataset. Protein candidates were selected from a list of protein hits that were detected in multiple replicates and minimally recovered with parental strain. Proteins known or predicted to interact with TgSKP1 are underlined. See Table S1 for origin of protein labels and more information about control data.

We next analyzed the TgFBXL2 interactome using co-immunoprecipitation (co-IP) methodology targeting the N-terminal 3xHA tag. MS-scale immunoprecipitation experiments were performed in 3 biological replicates, with 3 technical replicates each. We identified and quantified a total of 757 proteins: 317 at high confidence (FDR <1 %), 112 at medium confidence (1% < FDR < 5%), and 328 at low confidence (5% < FDR < 10%). Gene ontology analysis of the high and medium confidence interactors revealed that these genes encoded proteins that function in diverse biological processes with those associated with protein metabolism as the primary one identified **(Table S1).** Ten proteins satisfied the criterion of being detected by at least 2 peptides at an FDR rate of ≤1%, at a level ≥10-fold in ^HA^TgFBXL2 parasites vs. the untagged parental strain with a *P* <0.01 **(Fig. 1D)**. In addition to ^HA^TgFBXL2, these proteins include SCF components Cullin-1 and TgSKP1, several homologs of COP9-signalosome components (CSN1, CSN2, CSN3, CSN5, CSN6, and CSN7), a small ribonucleoprotein G homolog, and a small unknown protein comprising 185 amino acids and classified as a putative heat-shock protein 20 family member **(Table S1)**. Relaxing the 10-fold enrichment confidence to *P* ≤0.05 yields, in addition, a thioredoxin-like cytoplasmic protein, a filament type protein, and an abundant secretory protein not expected to contact TgFBXL2. The appearance of these likely false-positive hits suggests that the primary interactors accessible by this method have been achieved.

Cullin-1 is highly represented in the TgFBXL2 co-IPs **(Fig. 1E)** and has a similar number of amino acids but it cannot be assumed to be stoichiometrically associated with TgFBXL2 owing to the many variables involved in detecting and quantitating peptides that compose proteins. Nevertheless, it is clear that at least a substantial fraction of TgFBXL2 is associated with the SCF complex, which is supported by the presence of TgSKP1. The most striking observation is that 6 of the 8 predicted subunits of the COP9 signalosome (CSN) were detected at substantial levels. The CSN is an enzyme complex that mediates the deneddylation of Cullin-1, which in turn normally allows Cullin-1 to associate with CAND1. Cullin-1/CAND1 is unable to complex with FBP/SKP1 complexes. The stable binding of the CSN with SCF(TgFBXL2) complexes as implied by these findings is unusual and was not observed in an analysis of the interactome of TgFBXO1 (unpublished data). The findings suggest that a substantial fraction of the SCF(TgFBXL2) complex is held in an inactive state.

### Loss of TgFBXL2 Severely Impacts *Toxoplasma* Growth

To test whether TgFBXL2 is essential for tachyzoite growth, we first confirmed that ^HA^TgFBXL2 protein was successfully downregulated after 24 h treatment with 1 μg/mL ATC **(Fig. 2A)**. Using this ^HA^TgFBXL2 conditional mutant, we performed plaque growth assays and found that ^HA^TgFBXL2-depleted parasites were unable to form plaques, indicating that TgFBXL2 is essential for tachyzoite growth **(Fig. 2B).** Next, ^HA^TgFBXL2 and parental TATiΔKu80 parasites were grown for 24 h in the absence or presence of ATC and then fixed. Fixing after 24 h allows us to capture vacuoles before they lyse. Staining parasites by immunofluorescence (IFA) detection of the plasma membrane marker (SAG1), we first counted the numbers of vacuoles present to assess whether invasion was affected by loss of TgFBXL2. We found similar numbers of vacuoles were present **(Fig. 2C)**. Next, we examined replication by enumerating numbers of parasites per vacuole and found no significant differences **(Fig. 2D)**. These data indicate that TgFBXL2 did not play a significant role in either tachyzoite invasion or replication, which contrasts with the severe growth defect observed in the plaque assay.

**Figure 2.**
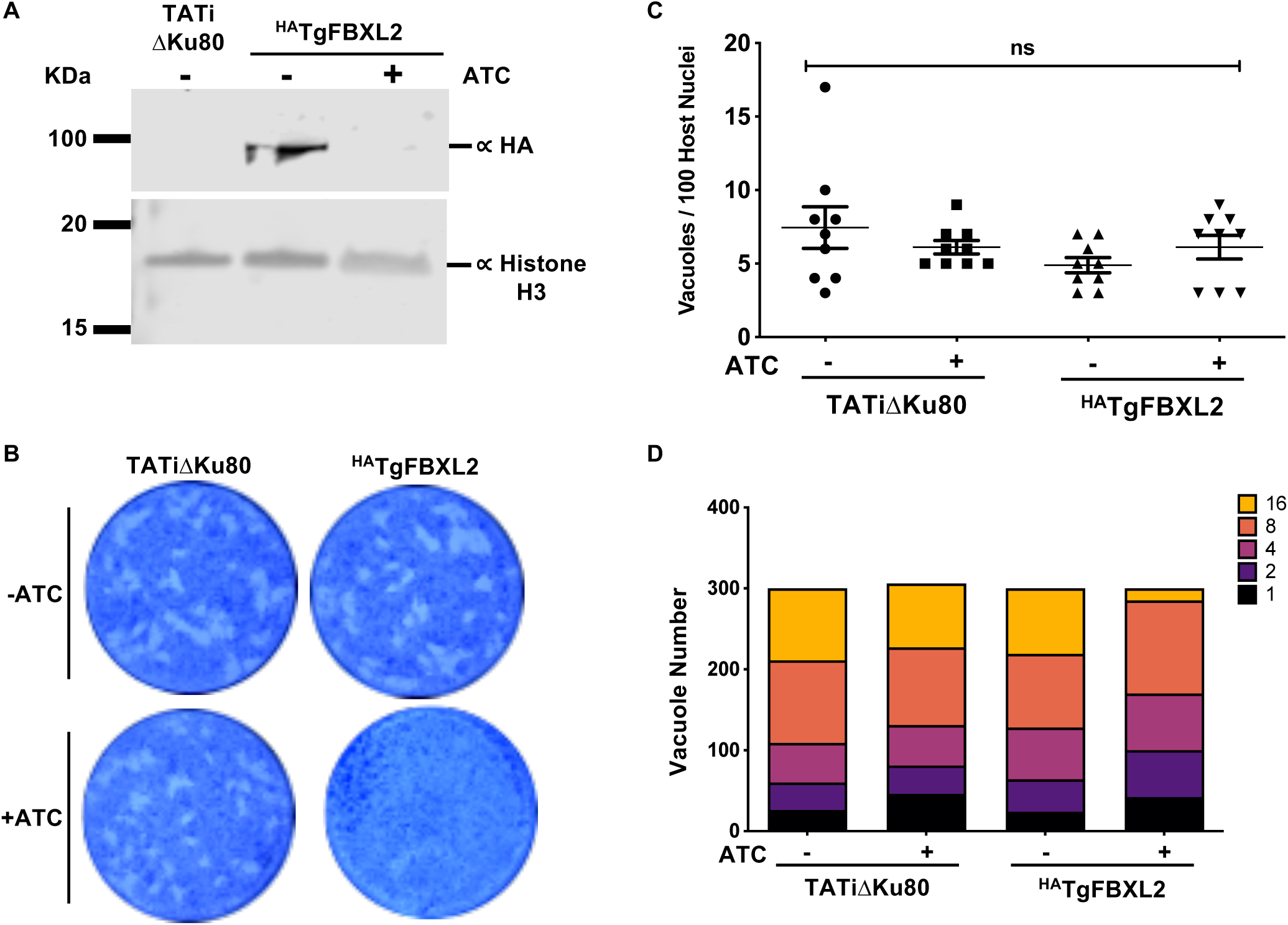
TgFBXL2 is Important for *Toxoplasma* growth. **(A).** Lysates from ^HA^TgFBXL2 or parental TATiΔKu80 tachyzoites grown for 24 h ± 1μg/mL ATC were Western blotted to detect ^HA^TgFBXL2 or Histone H3 as a loading control. **(B).** ^HA^TgFBXL2 or TATiΔKu80 parasites were grown for 7 days on HFF monolayers in the absence or presence of 1μg/ml ATC. Shown are representative images from 3 independent experiments performed in triplicate. (C and D). ^HA^TgFBXL2- or TATiΔKu80-infected HFF monolayers on coverslips were fixed after 24 h growth ± 1μg/mL ATC. **(C).** Invasion was assessed by determining numbers of vacuoles detected per 100 host cell nuclei. **(D).** Replication was determined by counting the number of parasites per vacuole. N = 3. Significance was analyzed using 2-way ANOVA.

### Loss of TgFBXL2 Affects Synchronicity of *Toxoplasma* Cell Cycle

Tachyzoites replicate by a unique process termed endodyogeny where the two daughter parasites develop within the mother [42, 43]. Normal endodyogeny progression depends on the tight regulation of centrosome duplication, which happens once and only once per cell cycle [44, 45]. To test whether loss of TgFBXL2 functions in centrosome duplication, ^HA^TgFBXL2 parasites were grown in the absence or presence of ATC, fixed 24 h later and stained with TgCentrin-1/IMC3 antibodies to detect the centrosome outer core and inner membrane complex (IMC), respectively. We found that although some ^HA^TgFBXL2-depleted parasites showed normal distribution of one centrosome per daughter cell **(Fig. 3A)** others showed an increased number of centrosomes that did not match the number of daughter cells **(* in Fig. 3A)**.

**Figure 3.**
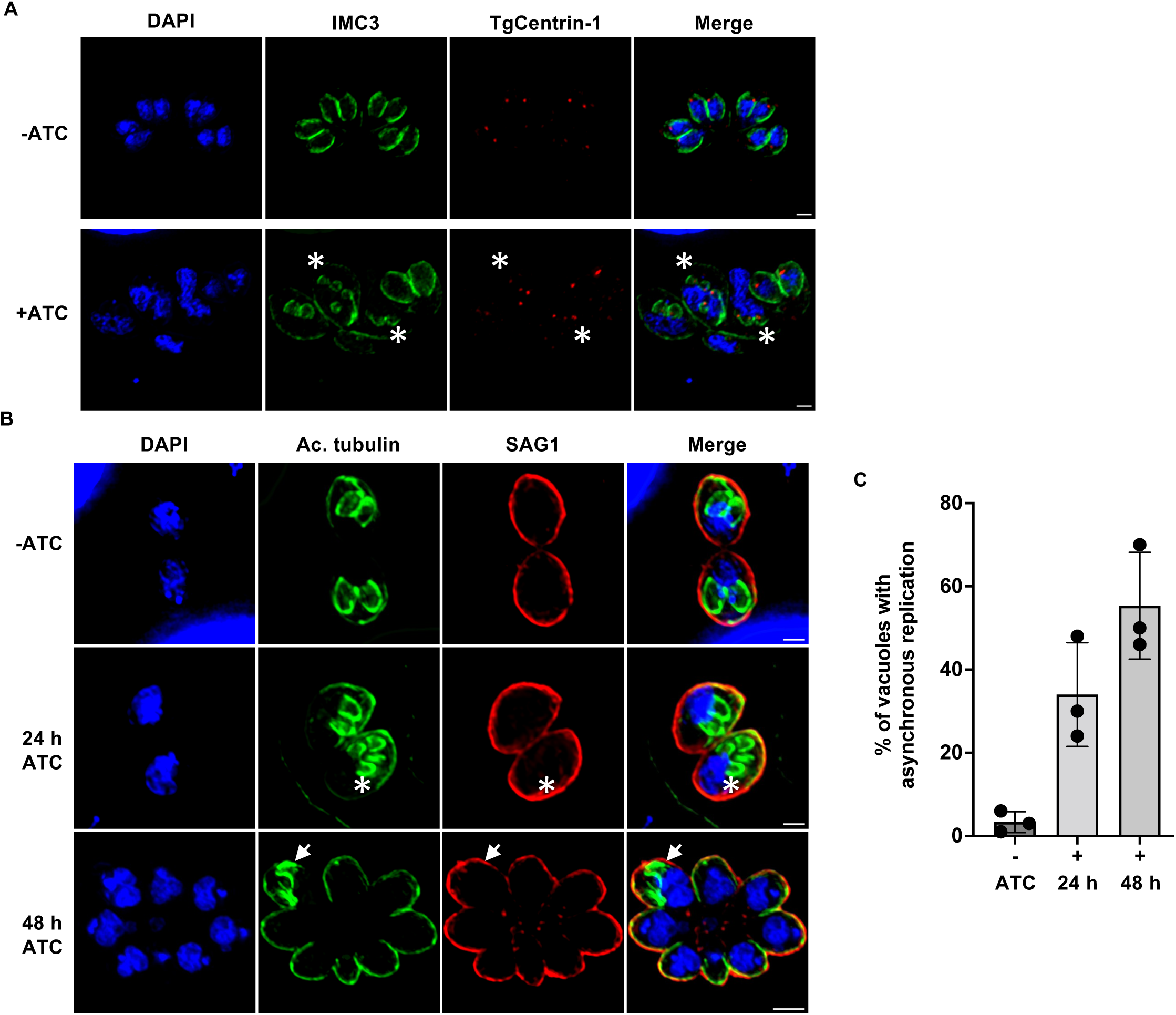
Loss of TgFBXL2 Affects *Toxoplasma* Cell Cycle Progression. **(A).** ^HA^TgFBXL2 parasites were grown for 24 h ± 1 μg/mL ATC. Cells were fixed and stained to detect IMC3, parasite’s centrosomes (TgCentrin-1) and DNA. (*) highlight ^HA^TgFBXL2-expressing parasites showing increased number of centrosome staining. **(B).** ^HA^TgFBXL2 parasites were grown for either 24 h or 48 h on HFF monolayers ± ATC, then fixed and stained to detect subpellicular microtubules (Ac. tubulin), plasma membrane (SAG1) and DNA. (*) highlight ^HA^TgFBXL2 parasites showing more than 2 daughter parasites. Bars = 1 µm in top and middle panel and 2 µm in bottom panel. **(C).** Quantification of vacuoles with asynchronous replication at the indicated times. Data represents averages ± standard deviations of 3 independent experiments with at least 50 parasites examined/experiment.

We next examined whether loss of TgFBXL2 negatively impacts daughter cell development. ^HA^TgFBXL2 parasites were grown in the absence or presence of ATC, fixed 24 or 48 h later and stained to detect SAG1 and acetylated tubulin, to mark recently synthesized subpellicular microtubules. In contrast to TgFBXL2-replete parasites that formed only two daughter cells per mother **(Fig. 3B upper panel)**, ^HA^TgFBXL2-depleted parasites showed an increased number of daughter cells per mother after 24 h ATC treatment **(* in Fig. 3B middle panel).** This phenotype became more prominent at 48 h ATC treatment **(Fig. 3C)** when it was also possible to observe parasites failing to divide **(* in Fig. 3B bottom panel)**. No defect in parasite replication was detected in the parental TATiΔKu80 strain growing in the presence of ATC (not shown). Collectively, these data suggest that TgFBXL2 is critical in controlling centrosome duplication and daughter cell development, which ultimately leads to asynchronized and unsuccessful divisions in ^HA^TgFBXL2-depleted parasites.

### Loss of TgFBXL2 Causes Apicoplast Biogenesis Defect

Loss of TgFBXL2 resulted in absence of detectable plaques even though they were still able to successfully divide, albeit at decreased efficiency. Plaque formation can also be abrogated when an essential chloroplast-like organelle named the apicoplast is either not functional or does not properly segregate into daughter parasites during endogeny [46, 47]. To test for apicoplast defects, ^HA^TgFBXL2 parasites were grown in absence or presence of ATC for 24 to 72 h and then stained to visualize the apicoplast (TgAtrx1), plasma membrane (SAG1), and DNA (DAPI). After 24 h ATC treatment, ^HA^TgFBXL2 depletion resulted in increased numbers of parasites lacking both TgAtrx1 staining and DAPI staining of the apicoplast genome **(Fig. 4A arrows)**. As time progressed, we noted increased numbers of TgAtrx1^-^ parasites **(Fig. 4B)**.

**Figure. 4.**
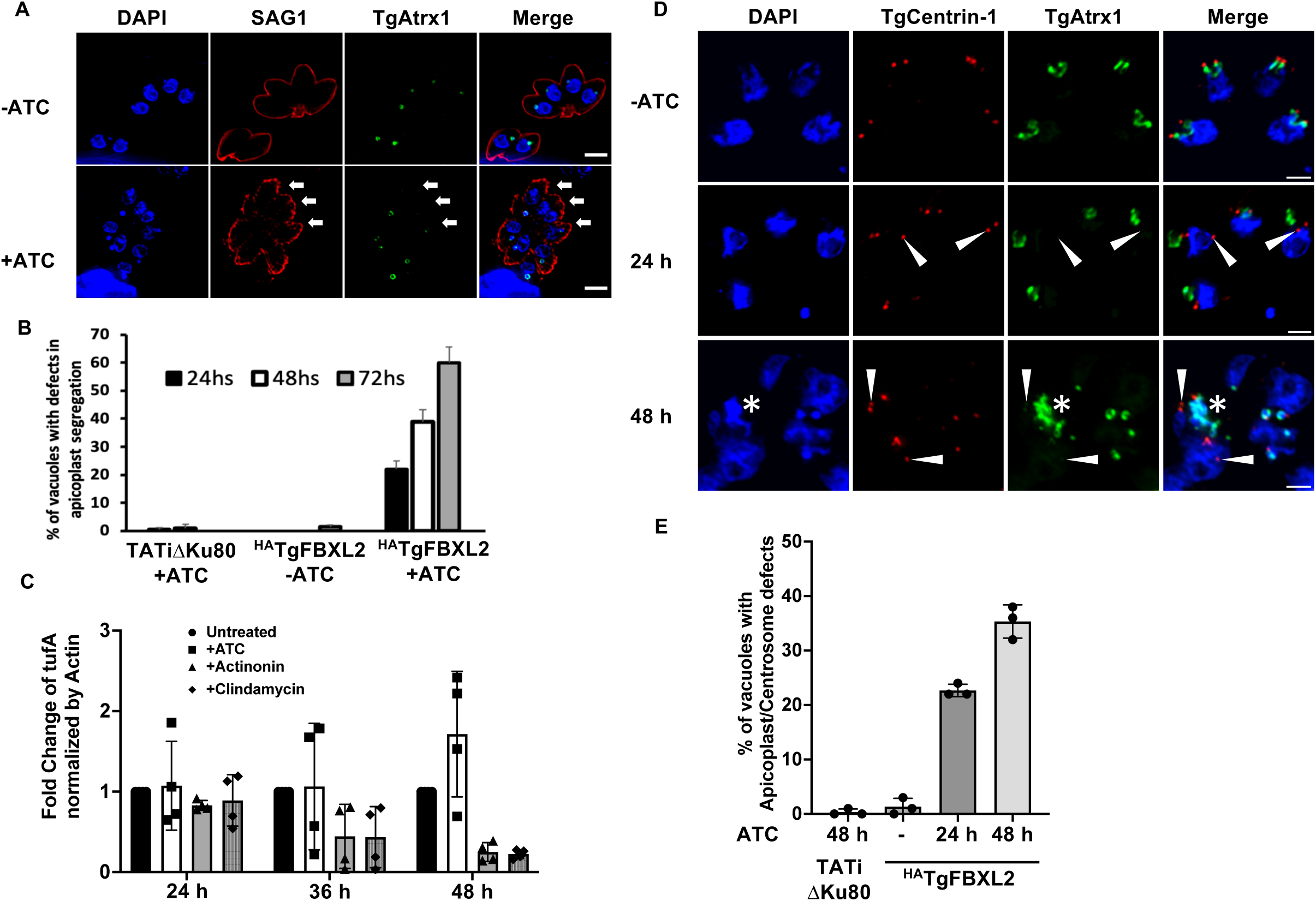
Loss of TgFBXL2 Causes Apicoplast Biogenesis Defect. **(A).** ^HA^TgFBXL2 parasites were grown for 24, 48, or 72 h on HFF monolayers ± 1 μg/mL ATC. Cells were fixed and stained to detect plasma membrane (SAG1), apicoplast (TgAtrx1) and DNA. Shown is representative image from parasites infected for 24 h. Arrows indicate ^HA^TgFBXL2 parasites lacking apicoplast staining. Bars = 2 µm. **(B).** Quantification of vacuoles with parasites showing apicoplast segregation defects at the indicated times. **(C).** qPCR was used to quantify apicoplast genome in ^HA^TgFBXL2 parasites grown for 24, 36, or 48 h on HFF monolayers ± ATC. Actinonin and clindamycin were used as positive inhibitors of apicoplast genome replication. Shown are means and standard deviations from 4 independent experiments. (P <0.05, one-way ANOVA). **(D).** ^HA^TgFBXL2 parasites were grown for either 24 h or 48 h on HFF monolayers ± ATC, then fixed and stained to detect parasite’s centrosomes (TgCentrin-1), apicoplast (TgAtrx1) and DNA. Arrowheads indicate ^HA^TgFBXL2 parasites in which TgCentrin-1 is not properly associated with apicoplast during cell division. Bars = 1 µm. **(E).** Quantification of vacuoles with parasites showing increased number of centrosomes that lack apicoplast interaction at the indicated times. Data represents averages ± standard deviations of 3 independent experiments with at least 50 parasites examined/experiment.

The apicoplast has a 35 Kb circular genome that duplicates prior to division of the apicoplast [48]. To test whether loss of TgFBXL2 affected apicoplast genome replication, ^HA^TgFBXL2 parasites were harvested from host cells after 24, 36, and 48 h ATC treatment, and genomic DNA was extracted for apicoplast genome quantification by qPCR. As controls, ^HA^TgFBXL2 parasites were also treated with either actinonin or clindamycin, both of which are known to inhibit apicoplast replication [49–51]. While ^HA^TgFBXL2 parasites treated with actinonin and clindamycin showed a significative decrease in apicoplast genomes, ^HA^TgFBXL2-depleted parasites did not **(Fig. 4C)**.

These data suggest that decreased TgFBXL2 expression affects apicoplast inheritance, which is dependent on tethering of the apicoplast and duplicated centrosomes. To test whether this process was impacted by loss of TgFBXL2, ^HA^TgFBXL2 parasites were grown in absence or presence of ATC for 24 and 48 h and then stained to visualize TgAtrx1, TgCentrin-1 and DAPI. ^HA^TgFBXL2 depletion resulted in increased numbers of centrosomes that did not interact with apicoplasts **(Fig. 4D, arrowheads)**. As time progressed, there was an increase in the number of vacuoles with parasites showing defects in apicoplast/centrosome interaction **(Fig. 4E)**. Moreover, at 48 h, apicoplasts failed to divide, becoming disorganized **(* in Fig. 4D)**.

### TgFBXL2 Localizes to a Specific Perinucleolar Region

To analyze TgFBXL2 localization ^HA^TgFBXL2 parasites and parental TATiΔKu80 were fixed and stained to detect ^HA^TgFBXL2, SAG1, and DAPI. ^HA^TgFBXL2 predominantly localizes to regions with low nuclear DAPI staining **(Fig. 5A).** Additionally, TgFBXL2 nuclear localization was proximal to, but did not overlap with, the nucleolus as evidenced by the SYTO RNA staining **(Fig. 5B)**.

**Figure 5.**
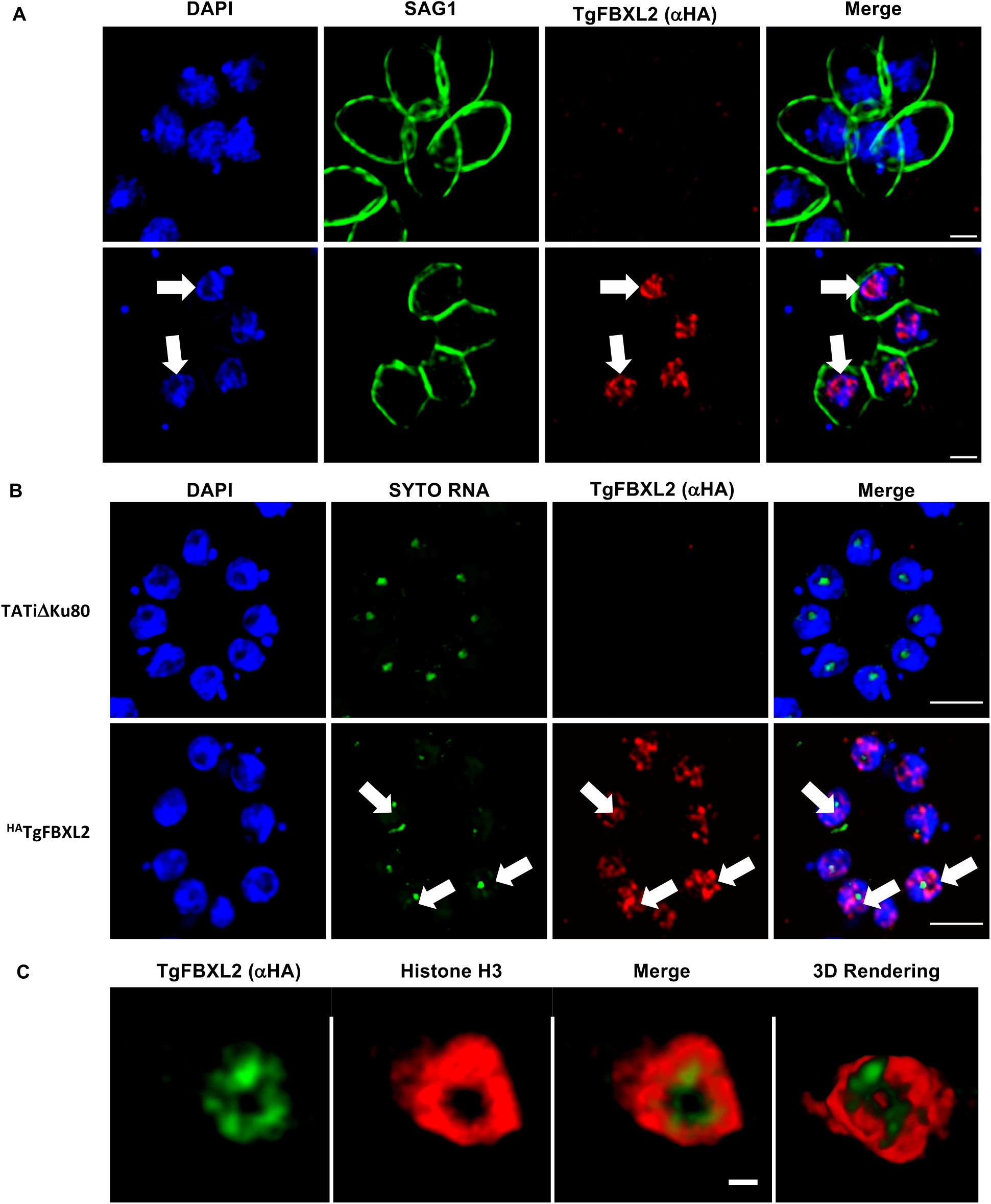
TgFBXL2 Localizes to a Perinucleolar Compartment. **(A).** ^HA^TgFBXL2-expressing parasites and parental TATiDKu80 were fixed and stained to detect ^HA^TgFBXL2 (aHA), plasma membrane (SAG1) and DNA. Arrows indicate HATgFBXL2 staining in areas staining weakly with DAPI. Bars = 1 µm. **(B).** ^HA^TgFBXL2 and parental TATiDKu80 tachyzoites were fixed and stained to detect ^HA^TgFBXL2 (αHA), nucleolus (SytoRNA) and DNA. Arrow highlights a parasite nucleus with ^HA^TgFBXL2 staining surrounding the nucleolus. Bars = 2 µm. **(C).** ^HA^TgFBXL2-expressing parasites were fixed and stained to detect ^HA^TgFBXL2 (αHA), and Histone H3 (αH3). Shown are still images from a movie of 3D rendering available as supplemental data. Bars = 0.5 µm.

Next, we used high-resolution <u>S</u>timulated <u>E</u>mission <u>D</u>epletion (STED) microscopy, to gain better insight into the distribution and organization of TgFBXL2 within the nucleus. Thus, ^HA^TgFBXL2 parasites were stained with anti-Histone H3 and anti-HA antibodies. ^HA^TgFBXL2 localization surrounded the central region where the nucleolus localizes, forming a ring that was proximal to, but distinct from, histone H3 **(Fig. 5C and Movies S1 and S2)**. Taken together, these data indicate that TgFBXL2 is an essential perinucleolar protein whose loss leads to defects in endodyogeny.

### Loss of TgFBXL2 Leads to Upregulation of Genes Involved in Sexual Commitment

TgFBXL2 nuclear localization and the loss of synchronicity in parasite replication pointed to a possible role in transcriptional regulation or chromatin remodeling. We therefore analyzed the transcriptomes by RNA-sequencing (RNAseq) of ^HA^TgFBXL2 parasites grown in the absence or presence of ATC for 24 and 48 h. ^HA^TgFBXL2 depletion led to an accumulation of 355 mRNAs at 48 h but not at 24 h **(Fig. 6A)**. No specific molecular or biological function was evidenced by GO analysis; however, GO analysis of cellular component revealed that 87% of up-regulated genes are membrane protein and the remaining genes encoded for cytoskeleton-associated proteins **(Fig. 6B)**.

**Figure 6.**
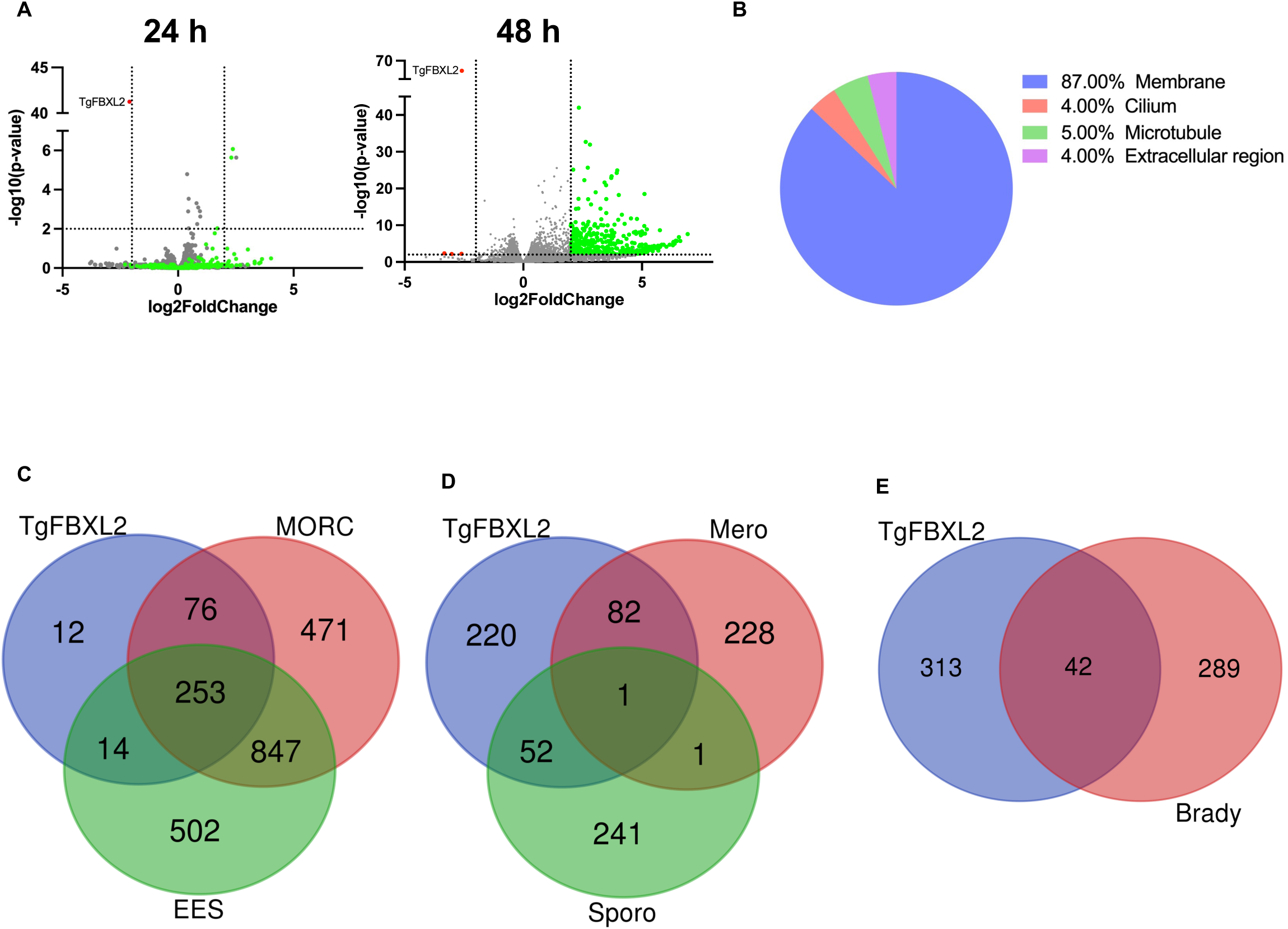
Loss of TgFBXL2 Upregulates Pre-Sexual Specific Genes Involved in Sexual Commitment. **(A).** Volcano plot displaying gene expression variations between the ^HA^TgFBXL2 parasites grown ± ATC for 24 h or 48 h (n=1549, Table S2). The red and green dots indicate those transcripts whose abundances are significantly down- and up-regulated genes, respectively, at 48 hpi, using adjusted *P* <0.01 (Bonferroni-corrected) and ± 2-fold change as the cut-off threshold. **(B).** Gene ontology for cellular component (CC) annotation of up-regulated genes in ^HA^TgFBXL2-depleted parasites. **(C).** Venn diagram comparing genes modulated by ^HA^TgFBXL2 to MORC or expressed by enteroepithelial stages parasites (EES). **(D & E).** Venn diagrams illustrating overlap between ^HA^TgFBXL2-upregulated genes (n = 355) and the RNAs expressed by merozoites, sporozoites, and bradyzoites.

Remarkably, about 92% of up-regulated genes in ^HA^TgFBXL2-depleted parasites overlap with genes induced by loss of a master regulator of *Toxoplasma* sexual development named MORC **(Fig. 6C)** [21]. Additionally, comparative RNA-seq analysis, using existing RNA seq data of enteroepithelial stage (EES) parasites [52], revealed that 75% of ^HA^TgFBXL2 up-regulated genes are specifically expressed in EES stages **(Fig. 6C)**.

Previous studies identified 312 genes exclusively expressed by merozoites [20, 53]. Interestingly, ^HA^TgFBXL2 depletion induced expression of merozoites-specific genes (n=83/355) and to a lesser extent, sporozoite-specific (n=53/355) **(Fig. 6D)** and bradyzoite-specific genes (n=42/355) **(Fig. 6E)** [53, 54]. This included genes that encode merozoite marker proteins such as GRA80 (TGME49_273980), GRA11B (TGME49_237800), MIC17c (TGME49_200230) and several merozoite-restricted surface proteins (SRS) **(Table 2)** [20, 55, 56]. Using IFA analyses, we confirmed that GRA80 protein **(Fig. 7A)** and ROP26, which can be expressed by both merozoites and bradyzoites **(Fig. 7B)** were detectable only after a 48 h ATC treatment.

**Figure 7.**
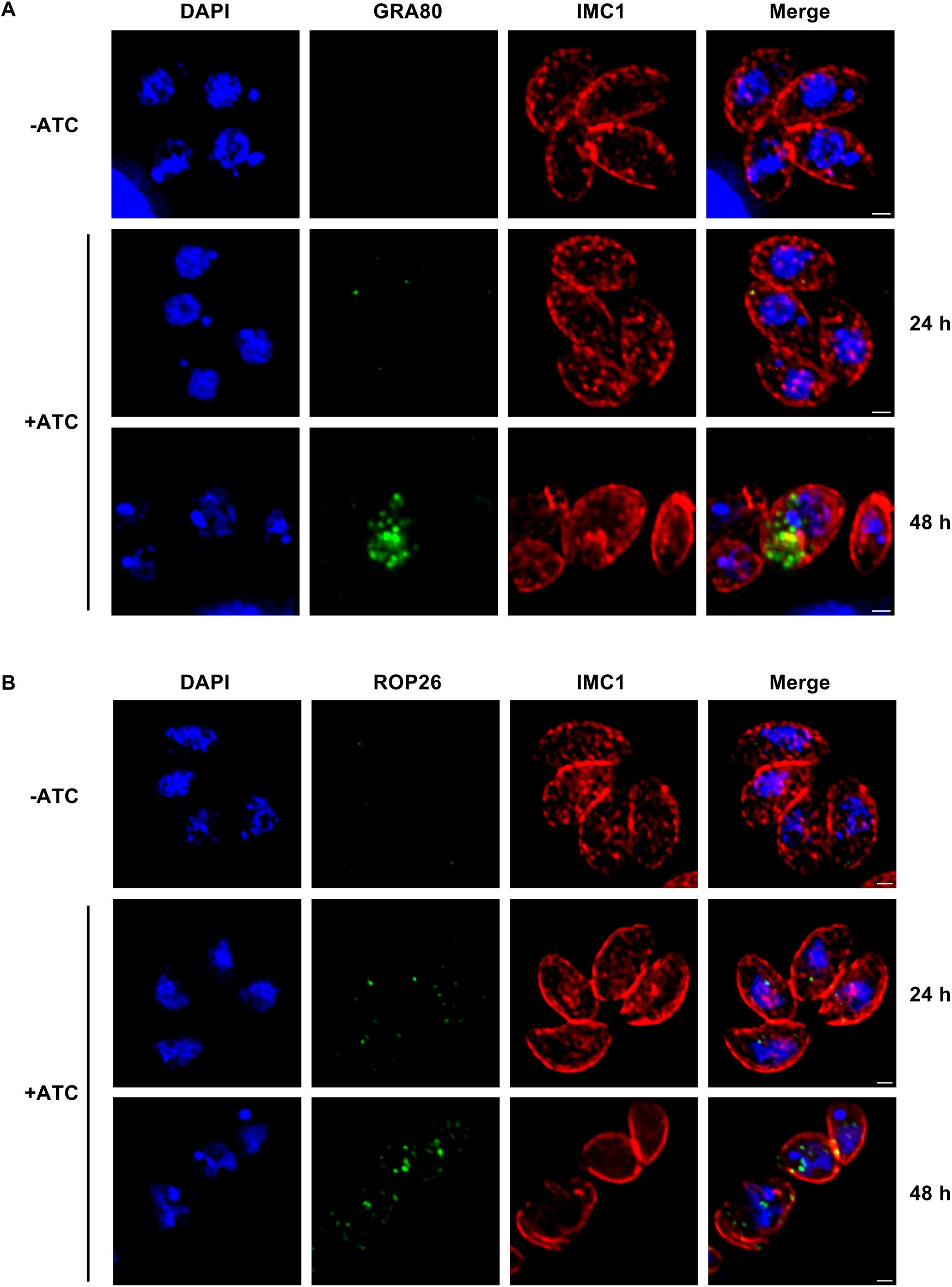
Merozoite-specific proteins Gra80 and ROP26 Are Expressed in TgFBXL2-Depleted Parasites. **(A).** ^HA^TgFBXL2 parasites grown for 24 h or 48 h ± 1 μg/mL ATC were fixed and stained to detect GRA80, IMC1 and DNA. (B). ^HA^TgFBXL2 parasites grown for 24 h or 48 h ± 1 μg/mL ATC were fixed and stained to detect ROP26, IMC1 and DNA. Bars = 1 µm.

### TgFBXL2 and MORC/HDAC3 Function Independently

MORC and HDAC3 function together in tachyzoites to repress the expression of other sexual stage–specific genes. Finding significant overlap between genes repressed by MORC/HDAC3 and TgFBXL2 raised the question of whether they act in unison or independently of one another. The TgFBXL2 and MORC/HDAC3 [21, 57] interactomes did not reveal direct interactions between them, and MORC was not detected by Western blotting of TgFBXL2 immunoprecipitates (not shown). To further assess potential interactions between the two, we used high-resolution STED microscopy to determine whether they colocalize. We found that although both localize to a peri-nucleolar region they do not overlap **(Fig. 8A & B).**

**Figure 8.**
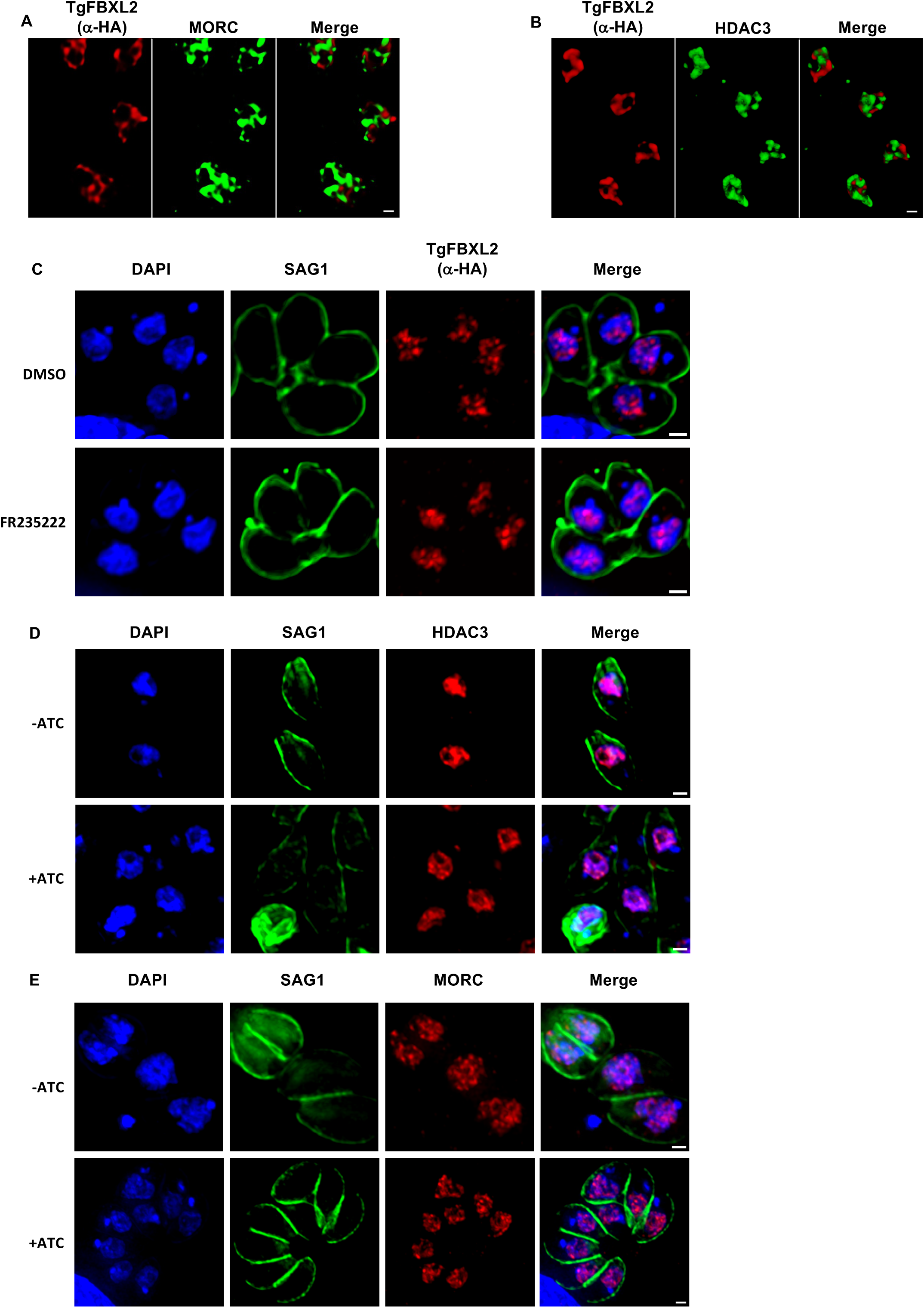
TgFBXL2 and MORC/HDAC3 Do Not Interact. **(A).** ^HA^TgFBXL2-expressing parasites were fixed and stained to detect ^HA^TgFBXL2 (αHA), and MORC. **(B).** ^HA^TgFBXL2-expressing parasites were fixed and stained to detect ^HA^TgFBXL2 (αHA) and HDAC3. Bars = 0.5 µm. **(C).** ^HA^TgFBXL2 parasites were grown for 24 h in the presence of either 100 nM FR235222 or DMSO as vehicle control. Cells were fixed and stained to detect ^HA^TgFBXL2 (αHA), SAG1 and DNA. **(D&E).** ^HA^TgFBXL2 parasites were grown for 48 h ± 1 μg/mL ATC. Cells were fixed and stained to detect DNA, SAG1 or either HDAC3 **(D)** or MORC **(E).**

Since MORC/HDAC3 bind chromatin, we tested whether TgFBLX2 did so as well by performing CHIP-Seq analysis of ^HA^TgFBXL2 and the parental untagged strain. The samples and input used as a control were purified, amplified, and subjected to next-generation sequencing. Reads were mapped to the *Toxoplasma* genome and normalized per million of mapped reads per sample. The data revealed no significant differences across all samples and replicates between the ^HA^TgFBXL2 tagged samples, the parental untagged samples, and the control tracks (**Fig S2**). To ensure that an inability to obtain sequence reads from the CHIPs did not reflect co-purifying a factor that inhibited the sequencing reactions, we used TapeStation analysis to test whether nucleic acid could be detected in the ^HA^TgFBXL2 IPs. Consistent with the CHiPSeq data, we failed to detect DNA in the CHIPs (**Fig S3)**. These data therefore indicate that TgFBXL2 does not directly interact with chromatin.

We next tested whether localization of either TgFBXL2 or MORC/HDAC3 was dependent on the other. ^HA^TgFBXL2 parasites were treated with FR235222, which is a HDAC3 inhibitor [58] and ^HA^TgFBXL2 localization was examined by IFA. We found that ^HA^TgFBXL2 perinucleolar localization was not impacted by the HDAC3 inhibitor **(Fig 8C)**. Similarly, HDAC3 and MORC localization was unaffected when ^HA^TgFBXL2 parasites were treated with ATC **(Fig 8D & E)**.

## DISCUSSION

FBPs regulate important roles in several biological functions [30, 31, 59–62]. Compared with the knowledge of metazoans, plants and fungi, our knowledge about FBPs in protozoan parasites is very limited. Previously, we identified 18 putative *Toxoplasma* FBPs and here describe the role of one of those FBPs, TgFBXL2 [40]. We confirmed the identification of TgFBXL2 as a FBP by demonstrating its interactions with core components of the SCF-E3 as well as multiple CSN subunits. TgFBXL2 was reported to be important for parasite fitness [39]; a finding we corroborate using a conditional expression system. Interestingly, the loss of TgFBXL2 had a modest replication delay, which contrasts with the loss of plaque formation when parasites were grown in the presence of ATC. This phenomenon was later explained by the loss of control of parasite replication as evidenced by development of multiple inviable daughter cells, a consequence of either organelle mis-segregation and/or expression of sexual-stage genes normally silenced in tachyzoites.

Our data suggest that a substantial fraction of TgFBXL2 is associated with the CSN. Highly conserved throughout eukaryotes, the CSN in plants, yeast, and humans consists of 8 canonical subunits [63, 64]. The corresponding *Toxoplasma* orthologs are predicted in Table S1, but it should be noted that homologs found in the eIF translation complex and the proteasome regulatory lid complex make the predictions ambiguous. However, the presence of six of the eight predicted *Toxoplasma* CSN subunits in the ^HA^TgFBXL2 interactome confirms their identity as CSN subunits. The other two subunits, CSN4 and CSN8, were detected in the co-IPs but were not enriched over the un-tagged strain control. This alone does not exclude their presence in the CSN complex, but additional co-immunoprecipitation assays will be required to determine whether they are *bona fide* CSN components. Interestingly, CSN4 and CSN8 were also not detected in a study of the CSN in *Entamoeba histolytica* [65], whereas all 8 were detected in another protist, the social amoeba *Dictyostelium* [66]. Structural studies in other organisms reveal that the CSN covers a large fraction of one face of the SCF, thereby interfering interactions with ubiquitin E2 ligase and SCF target substrate [67]. The interactome data are also supported by a failure to detect the small ubiquitin like protein NEDD8 in the TgFBXL2 interactome since CSN5 is a deNEDDylase [64, 68]. Why the SCF(TgFBXL2) is in a stable inactive complex with CSN is unclear, but studies in other organisms show that inositol hexakisphosphate is a metabolic regulator of the complex [66]. Thus, as a major SCF complex, SCF(TgFBXL2) appears poised to act in substrate degradation but presumably dependent on an unknown trigger to release CSN inhibition. It is also possible that the CSN functions independently of the SCF as it does in plants regulating seed germination [69]

Decreased TgFBXL2 expression had severe effects on parasite growth and gene expression. In the absence of TgFBXL2, *Toxoplasma* tachyzoites became unviable due to increases in the numbers of parasites that either divided asynchronously or lacked an apicoplast or nucleus. We did not examine other organelles but would expect similar defects in Golgi segregation as it also associates with duplicated centrosomes during endodyogeny [70, 71]. Loss of TgFBXL2 also severely affected parasite gene expression with the upregulation of genes primarily associated with pre-sexual and sexual stage development. Currently, we are unable to determine whether these growth and gene expression phenotypes are linked or are due to TgFBXL2 functioning in two distinct processes, perhaps due to multiple substrates as typical for many FBPs [72]. We believe that the former is more likely, because TgFBXL2 was only detectable within the nucleus as are other factors known to impact merozoite gene expression such as MORC/HDAC3 and the Api2XI-2 and Api2XII-1 transcription factors [21, 73]. Thus, we propose that the growth defects observed in TgFBXL2-depleted parasites is due to aberrant expression cell cycle proteins whose activities cannot be controlled by tachyzoite-encoded checkpoint proteins.

MORC and HDAC3 repress the expression of merozoite genes and are recruited to the promoters of these genes by AP2XI-2 and AP2XII-1 transcription factors [73]. These two transcription factors regulate the majority of the MORC/HDAC3 target genes and although most target genes are regulated by both, some only require depletion of one or the other. We did not find any specific enrichment of TgFBXL2-repressed genes in either category and together with a lack of biochemical or functional interactions between them, these data point to a model in which TgFBXL2 and MORC/HDAC3 function independently of one another. Indeed, immunofluorescence analysis of ^HA^TgFBXL2 parasites treated with the HDAC3 inhibitor did not reveal significant changes in TgFBXL2 localization or abundance. Thus, how TgFBXL2 regulates gene expression remains enigmatic. One model is that TgFBXL2 promotes the turnover of proteins that activate merozoite gene expression, and these genes are likely to not include MORC, HDAC3, AP2IX-2, and AP2XII-1. By acting at the level of the proteome, TgFBXL2 may serve as a back-up to transcriptional regulation to guard against spurious merozoite gene expression and protect the current state of differentiation. It also remains unclear how TgFBXL2-dependent protein ubiquitination is inhibited to allow expression of the genes that it represses. Our prediction is that that alterations in post-translational modifications of TgFBXL2 substrates likely allows for differences in ubiquitination and future work will identify these proteins and modifications.

## MATERIALS AND METHODS

### Cell lines and *Toxoplasma* strains

*Toxoplasma* strain TATi-RHΔku80 [41] was cultured in human foreskin fibroblasts in Dulbecco’s Modification of Eagle’s Medium (DMEM) (VWR; Radnor, PA, USA) supplemented with 10% fetal bovine serum (VWR), 2 mM L-glutamine (VWR), and 100 IU/mL penicillin – 100 µg/mL streptomycin (VWR). Parasites were released from host cells by passage through a 27-gauge needle [74]. All parasite strains and host cell lines were routinely tested for mycoplasma contamination with the MycoAlert Mycoplasma Detection Kit (Lonza, Basel, Switzerland) and found to be negative.

### Generation TgFBXL2 Constructs

Using the ToxoDB gene model for TgGT1_313200 as a guide, a ^HA^TgFBXL2 conditional expression mutant containing an anhydrotetracycline (ATC) responsive promoter derived from gene promoter for SAG4 and an N-terminal 3x-HA tag in frame with TgFBXL2 start codon was generated. The 5’ end of TgFBXL2 was amplified by using the following primers (regions of homology to TgFBXL2 are underlined and vector annealing sequences are in lower case) forward 5′-atgttccagattatgcc<u>ATGCTGGAGGCGAGGAAC</u>-3′ and reverse 5′-cgcggtggcggccgc<u>ACGGTGAAAAGAGGAGAAGC</u>-3′. The fragment was cloned by Gibson Assembly (New England Biolabs; Ipswich, MA, USA) into ptetO7sag4-HA-CEP250-DHFR-TS, replacing the CEP250 cassette [75]. The resulting construct was linearized by SgrDI, transfected into the TATiΔku80 strain [41, 75], and clones isolated by limiting dilution using pyrimethamine resistance.

### Invasion, Replication and Plaque Assays

For all assays, parasites harvested from *Toxoplasma*-infected HFF monolayers were released by syringe lysis by passage through a 27-gauge needle followed by washing in serum-free medium. Synchronized invasion, replication, and plaquing assays were performed as described previously [40].

### Immunoprecipitation

^HA^TgFBXL2 tachyzoites were harvested by syringe lysis and washed in ice-cold PBS. Parasites (1*10^8^) were resuspended in IP buffer containing 50 mM Tris-HCl (pH 7.4) with 1% Triton X-100, 100 mM NaCl, 1 mM NaF, 0.5 mM EDTA, 0.2 mM Na_3_VO_4_, 1X protease inhibitor cocktail (Thermo Fisher Scientific. Waltham, MA), incubated on ice for 30 min and then subjected to three pulses of sonication for 30 seconds and 25% amplitude each. Lysates were clarified by centrifugation at 16,000 × g, incubated with mouse anti-HA mAb clone 12CA5 conjugated with protein G agarose beads (Sigma-Aldrich) for 16 h at 4°C, and sequentially washed three times with IP buffer by centrifugation at 3500 x g for 5 min. Immune complexes were separated with SDS-PAGE and then Western blotted with rat anti-HA antibody (clone 3F10, Roche), or rabbit anti-TgSkp1 affinity purified polyclonal antibody UOK75 [76],

### Large scale immunoprecipitation for MS analysis

^HA^TgFBXL2 and parental strains parasites (2.5*10^8^) harvested as described above were resuspended in 1 µL Resuspension Buffer (50 mM HEPES (pH 7.4) with 0.5% NP-40, 100 mM NaCl, supplemented with 10 µg/mL aprotinin, 10 µg/mL leupeptin, 1 mM PMSF, 1 mM NaF, and 0.2 mM Na_3_VO_4_), and incubated on ice for 5 min. Lysates were clarified by centrifugation at 21,000 × g, at 4°C and incubated with mouse anti-HA mAb clone 12CA5 conjugated and crosslinked to protein A/G magnetic agarose beads (Pierce; Rockford, IL. Thermo Fisher Scientific. Waltham, MA, USA). After rotation at 4°C for 1 h, beads were magnetically captured, and the supernatant removed. The beads were successively washed three times in Resuspension Buffer without protease inhibitors, three times in 10 mM Tris-HCl (pH 7.4), 50 mM NaCl and once in 50 mM NaCl. Proteins were eluted off the beads by gentle rocking for 15 min at 22°C in 133 mM triethylamine (TEA, Sequencing Grade, Pierce), immediately neutralized with acetic acid, and dried in a vacuum centrifuge. Samples were then reduced, alkylated, and converted to peptides with trypsin; peptides were then captured on C18 Zip-tips and released for MS analysis, all essentially as described [77].

### Proteomic analysis

Peptides were separated on a C18 nano-column (PepMap 100 C18 series. Thermo Fisher Scientific. Waltham, MA, USA) using an Ultimate 3000 nano-HPLC, and directly infused into a Q-Exactive Plus Orbitrap Mass Spectrometer (Thermo Fisher), as previously described [77]. Raw files were then processed in Proteome Discoverer 2.5 using a *Toxoplasma* GT1 protein database containing 8,450 unique proteins (UniProt Proteome ID UP000005641), modified to include a list of 179 common ectopic contaminants [78] essentially as described [77]. Proteome Discoverer 2.5 calculated protein abundances for proteins of high (FDR<0.01) and medium confidence (FDR<0.05), and identified with 2 or more peptides, were compared between controls (parental strain) and samples (^HA^TgFBXL2 strain) in MetaboAnalyst 5.0 data analysis tool [79]. Proteins enriched >10-fold with a *P* <0.05 in ^HA^TgFBXL2 parasites vs. the untagged parental strain, were considered significant interactors of TgFBXL2. The MS proteomics data (listed in Table S1) are deposited in the ProteomeXchange Consortium via the PRIDE [80] partner repository with the dataset identifier PXD046583 and 10.6019/PXD046583..

### Western Blotting

Parasites were pelleted by centrifugation at 2000 × g for 8 min at 4°C and lysed in boiling SDS-PAGE sample buffer containing 2-β mercaptoethanol. Equivalent protein amounts of lysates were separated by SDS-PAGE gels, transferred to nitrocellulose membranes, and blocked with Odyssey Blocking Buffer (LI-COR Biosciences, Lincoln, NE), blotted using appropriate primaries antibodies. Blots were imaged using a LI-COR Odyssey scanner and analyzed using Image Studio software (LI-COR; Omaha, NE, USA).

### Immunofluorescence Microscopy

*Toxoplasma*-infected HFFs grown on coverslips were fixed with 4% w/v paraformaldehyde in phosphate-buffered saline (PBS) for 20 min at room temperature. Cells were permeabilized with 0.1% Triton X-100 in PBS for 10 min, blocked in 5% w/v bovine serum albumin (BSA) in PBS for 60 min, incubated overnight with primary antibodies at 4°C, and then incubated for 60 min with Alexa Fluor 488- or Alexa Fluor 594-conjugated antibodies (1:2000, Thermo Fisher Scientific). DNA was stained by incubation with 1 μg/mL DAPI (Thermo Fisher Scientific) for 5 min followed by mounting in VECTSHIELD medium (Vector Labs; Burlingame, CA, USA). Images were acquired using a 100X Plan Apo oil immersion 1.46 numerical aperture lens on a motorized Zeiss Axioimager M2 microscope equipped with an Orca ER charge-coupled-device (CCD) camera (Hamamatsu, Bridgewater, NJ). Images were collected as a 0.2-μm z-increment serial image stacks, processed using the Volocity, version 6.1, Acquisition Module (Improvision Inc., Lexington, MA). Images were deconvolved by a constrained iterative algorithm, pseudo colored, and merged using the Volocity Restoration Module. All images are maximal projection images and were processed similarly. All data were quantified from at least 50 randomly selected images for each condition from three independently performed experiments.

For STED microscopy, *Toxoplasma*-infected HFFs grown on coverslips were treated as mentioned above, but the samples were incubated with both Abberior Star Orange, goat anti-mouse IgG and Abberior Star Red, goat anti rabbit IgG (Abberior, Göttingen, Germany) secondary antibodies, followed by mounting in ProLong Glass Antifade (Thermo Fisher Scientific). Images were acquired as 0.15 μM z-increment serial images using Leica Hyvolution Acquisition Software (Leica; Buffalo Grove, IL) with a Leica TCS SP8 confocal microscope equipped with a 100x/1.47 TIRF oil immersion objective lens and both white light laser (470 nm-670 nm) and 405 nm diode laser. Image datasets were then deconvolved and 3D volume was generated by Leica visualization software. Images from same experiments were processed using identical settings.

### Illumina library preparation and RNA sequencing

^HA^TgFBXL2 parasites were grown in the presence of ATC for either 24 or 48 h. ^HA^TgFBXL2 parasites growing in the absence of ATC were used as control. After ATC treatment, parasites were harvested from infected HFF monolayers by scraping and passage through a 27-gauge needle, centrifuged at 2000 × g for 8 min at room temperature, resuspended in phosphate-buffered saline, counted, pelleted and total RNA was extracted using SV Total RNA Isolation System (Promega; Madison, WI).

Agilent 2100 Bioanalyzer was used to determine the integrity, purity, and concentration of RNA samples. RNA integrity (RIN) score of 6.5 or above was considered acceptable for further analysis. Total RNA was enriched for mRNA using poly-(A)-selection (Illumina; San Diego, CA). NEB stranded RNA library prep kit (NEB) and NEB Ultra II RNA library prep kit (NEB) were used to prepare complementary DNA (cDNA) libraries for all ^HA^TgFBXL2 samples, according to manufacturer’s protocol. RNA sequencing was carried out on an Illumina HiSeq2500 (Illumina) with a mid-output 75-cycle paired end with 10-20 million reads per sample at the Genomics and Bioinformatics core facility at the University at Buffalo.

### Differential gene expression analysis

Per-cycle basecall (BCL) files generated by the Illumina HiSeq2500 were converted to per-read FASTQ files using bcl2fastq version 2.20.0.422 with default settings. FastQC version 0.11.5 was used to review the sequencing quality while FastQ Screen version 0.11.1 was used to determine any potential contamination. FastQC and FastQ Screen quality reports were summarized using MultiQC version 1.5 [81]. Genomic alignments were performed using HISAT2 version 2.1.0 using default parameters [82]. To differentiate between bacterial vs *Toxoplasma* RNA, the resulting reads were aligned to genome annotation of GT1 strain available at https://toxodb.org/toxo/app. MultiQC software was used to summarize alignment as well as feature assignment statistics [81]. Differentially expressed genes were detected using the Bioconductor package DESeq2 version 1.20.0 [83]. Genes with one count or less were filtered out, and alpha was set to 0.05. Log2 fold-changes were calculated using DESeq2 using a negative binomial generalized linear models, dispersion estimates, and logarithmic fold changes integrated with Benjamini-Hochberg procedure to control the false discovery rate (FDR). A list of differentially expressed genes (DEGs) was generated through DESeq2. We defined a significant up or downregulation as a fold change ≥2 with FDR value <0.05. The PCA plots were generated in ggplot2 package, and the Venn diagram plots were made using the GraphPad Prism (GraphPad, La Jolla, CA).

### ChIP Seq

Approximately 3×10^8^ untagged and ^HA^TgFBXL2 tagged parasites each were pelleted and crosslinked with formaldehyde (1.6%), then quenched with glycerine ( to final concentrations of 0.25 mM) and followed by a series of washes with PBS. The resulting pellet was resuspended in 1 mL nuclear extraction buffer (10 mM HEPES, 10 mM KCl, 0.1 mM EDTA, 0.1 mM EGTA, 1 mM DTT, 0.5 mM AEBSF, 1X Roche protease inhibitor, 1X Roche phosphatase inhibitor) followed by a 30’incubation on ice. 10% Igepal CA-630 (final concentration 0.25%) was added to each sample followed by passage through a 26G needle and centrifuged at 5,000 rpm to obtain the nuclear pellet. Nuclear pellets were resuspended in shearing buffer (0.1% SDS, 1 mM EDTA, 10 mM Tris-HCl pH 7.5, 1X Roche protease inhibitor, and 1X Roche phosphatase inhibitor) and transferred into 130 μl Covaris tubes (Covaris; Woburn, MA). Samples were then sonicated using a Covaris S220 sonciator (under the following settings: 5 min, Duty cycle 5%, Intensity 140 W, 200 Cycles/burst, 4°C) before adding equal volumes of ChIP dilution buffer (30 mM Tris-HCl pH 8, 3 mM EDTA, 0.1% SDS, 30 mM NaCl, 1.8% Triton X-100, 1X protease inhibitor, 1X phosphatase inhibitor). Samples were centrifuged at 13,000 rpm for 10 min at 4°C. For each sample, 13 μl protein A agarose/salmon sperm DNA beads were washed 3 times with ChIP dilution buffer without inhibitors. The washed beads were added to the diluted chromatin for 1 hr at 4°C with agitation to pre-clear the samples. ~10% of each sample by volume was set aside as input and 2 μL of antibody HA (Abcam ab9110) was added to the remaining sample and incubated overnight at 4°C with rotation. To each sample, 25 μL of washed protein A agarose/salmon sperm DNA beads with ChIP buffer were blocked with 1 mg/ml BSA for 1 hr at 4°C, re-washed, and added to each sample for 1 hr rotation at 4°C. The bead/antibody/protein complexes were washed a total of 8 times with 15’ intervals per wash): twice with low salt buffer (1% SDS,1% Triton X-100, 2 mM EDTA, 20 mM Tris-HCl pH 8, 150 mM NaCl), twice with high salt buffer (1% SDS,1% Triton X-100, 2 mM EDTA, 20 mM Tris-HCl pH 8, 500 mM NaCl), twice with LiCl buffer (0.25 M LiCl, 1% NP-40, 1% Na-deoxycholate,1 mM EDTA, 10 mM Tris-HCl, pH 8.1), and twice with TE (10 mM Tris-HCl pH 8, 1 mM EDTA) buffer. DNA was then eluted from the beads with two 250 μL washes of elution buffer (1% SDS, 0.1M sodium bicarbonate) followed by the addition of NaCl (55ul of 5M) to reverse crosslink overnight at 45°C. RNAse A (15 μL of 20 mg/mL) was added and incubated at 37°C for 30;. Following this, they underwent proteinase K (2 μL 20 mg/mL) digestion at 45°C for 2 hours. The DNA was phenol/chloroform extracted and then ethanol precipitated overnight. After precipitation, the samples were centrifuged at 13,000 rpm for 30 min at 4°C, forming pelleted DNA, washed with 80% ethanol, re-pelleted, and resuspended in 50 μL of nuclease-free water. The DNA was purified with AMPure XP beads and prepared using a KAPA Hyperprep kit (KK8504), for sequencing by the NextSeq 500 sequencing platform (Illumina).

ChIP-seq read quality was analyzed using FastQC (https://www.bioinformatics.babraham.ac.uk/projects/fastqc/) and adapters and low-quality bases trimmed using Trimmomatic (http://www.usadellab.org/cms/?page=trimmomatic) and Sickle (https://github.com/najoshi/ sickle). Reads were then mapped against the ToxoDB-62_TgondiiME49 assembly using Bowtie2 (v2.4.4) [84], with uniquely mapped fragments and correctly paired reads using Samtools (v1.11) (http://samtools.sourceforge.net). Then, PCR duplicates were removed with PicardTools MarkDuplicates (v2.18.0)(Broad Institute). To obtain per nucleotide coverage and generate browser tracks, BedTools (v2.27.1) and custom scripts were used, normalizing counts to millions of mapped reads. Chromosome tracks were viewed using IGV (Broad Institute), then visualized using a custom script.

### TapeStation Analysis

Tachyzoite-infected monolayers (5×10^8^ parasites) were crosslinked in 1% formaldehyde, quenched with 125 mM glycine and rinsed in cold PBS before scraping in cold PBS and syringe-lysing to release intracellular parasites. After washing in cold PBS, parasite pellets were resuspended in cytoplasmic lysis buffer (85mM KCl, 5mM HEPES pH 8.0, 0.5% NP-40, 1mM phenylmethylsulfonyl fluoride (PMSF), cOmplete Protease Inhibitor Cocktail (Roche)) and incubated on ice for 10 min. The nuclear pellets were resuspended in nuclear lysis buffer (50mM Tris-HCl pH 8.0, 10mM EDTA, 1% SDS, 1mM PMSF, cOmplete Protease Inhibitor Cocktail (Roche)) and vortexed at 4°C for 30 min. Nuclear extracts were sonicated at 4°C with a Q-Sonica Q800R3 sonicator (Qsonica, Newtown, CT) at 75% amplitude for 12.5 minutes in intervals of 30 sec pulse on and 30 sec pulse off. Insoluble cell debris was cleared from the sample by centrifugation and soluble fraction was diluted 10-fold in IP dilution buffer (167mM NaCl, 16.7mM Tris-HCl pH 8.0, 1.2mM EDTA, 1.10% Triton X-100, 0.01% SDS, 1mM PMSF, cOmplete Protease Inhibitor Cocktail). The sample was precleared with Protein G magnetic beads (Pierce) for 1 hour at 4°C and incubated with 25 μl HA magnetic beads (Pierce) overnight at 4°C with rocking. For ChIP of H3K9ac, samples were incubated with 3 µg of rabbit anti-H3K9ac antibody (Active Motif; Carlsbad, CA) overnight at 4°C with rocking, followed by recovery with 25 µl Protein G magnetics beads for 3 h at 4°C. Beads were washed three times with Low Salt (150_mM NaCl, 20_mM Tris-HCl, 2_mM EDTA, 1% Triton X-100, 0.1% SDS), High Salt (500_mM NaCl, 20_mM Tris-HCl, 2_mM EDTA, 1% Triton X-100, 0.1% SDS) and LiCL (0.25_M LiCl, 10_mM Tris-HCl, 1_mM EDTA, 1% NP40, 1% deoxycholate) wash buffers, followed by two washes in TE buffer. Samples were eluted from beads in 1% SDS/TE buffer pH 8.0 and incubated overnight at 65°C to reverse crosslinks. Eluted DNA was recovered by phenol-chloroform extraction and sodium acetate precipitation with glycogen. DNA pellets were resuspended in TE buffer for downstream quantification and analysis by Qubit Fluorometer (ThermoFisher) and TapeStation 4200 automated electrophoresis (Agilent; Santa Clara, CA).

### Statistical analyses

Data were analyzed by one-way ANOVA with Tukey’s post hoc test or Student’s t test performed with GraphPad Prism (GraphPad, La Jolla, CA).

## FIGURE LEGENDS

**Figure S1. (A).** Predicted domain organization after HA-tagging. LRR = leucine-rich repeat, NLS = monopartite nuclear localization sequence. LRR3 and LRR4 are predicted by alphafold (see panel D), but are not identified by sequence prediction algorithms. Regions not labeled are poorly conserved. **(B).** Predicted amino acid sequence (from Toxo.db). Predicted domains are labeled and numbered as indicated. Underlined sequences were confirmed by mass spectrometry. Double underlined sequence represents the N-terminal 3X HA-tag. **(C).** Organization of amino acids 305-832 as predicted by alphaFold-2 (https://alphafold.ebi.ac.uk/entry/S8EUA4).

**Figure S2.** ChIP-Seq results visualization of HA tagged TgFBXL2 parasite line (TgFBXL2-HA) across all 13 of *Toxoplasma*’s chromosomes. Reads were normalized to millions of mapped reads. Tracks correspond to TgFBXL2-HA tagged parasites in blue, wild type parasites in red, and an input control in black. Former chromosomes 7b and 8 were combined into the recently suggested chromosome 13 [85, 86].

**Figure S3.** TgFBXL2 is not chromatin associated. Tapestation analysis of DNA purified from ^HA^TgFBXL2 ChIP. Gel image representing three ChIP DNA samples from ^HA^TgFBLX2 immunoprecipitated with anti-HA, ^HA^TgFBLX2 immunoprecipitated with anti-H3K9ac (positive control) and the parental TATi parasites immunoprecipitated with anti- HA. Electropherograms from each sample depict marker peaks at 15 and 10,000 bp, with a peak of immunoprecipitated DNA at 195 bp present only in the positive control.

**Table S1. TgFBXL2 Interactome Analysis.**

**Table S2: List of Genes Differentially Expressed by TgFBXL2-Depleted Parasites**

## ACKNOWLEDGEMENTS

We thank Drs. Marc Jan Gubbels, Elena Suvorova, and Zhicheng Dou for providing reagents and Dr. Adane Nigatu, Hubbard Center for Genome Studies, UNH for assistance with Qubit and TapeStation analyses. This work was supported by NIH Grant 1R01 AI-150240 to IJB and CMW; R01AI136511 aR01 AI158417 to KLR. VJ was supported as a Project Lead by the Center of Integrated Biomedical and Bioengineering Research through a grant from the National Institute of General Medical Sciences of the National Institutes of Health under Award Number P20GM113131. The funders had no role in study design, data collection and interpretation, or the decision to submit the work for publication.

## REFERENCES

1. Pappas G, Roussos N, Falagas ME. Toxoplasmosis snapshots: global status of *Toxoplasma gondii* seroprevalence and implications for pregnancy and congenital toxoplasmosis. Int J Parasitol. 2009;39(12):1385–94. Epub 2009/05/13. doi: 10.1016/j.ijpara.2009.04.003. PubMed PMID: 19433092.

2. Dubey JP. History of the discovery of the life cycle of *Toxoplasma gondii*. Int J Parasitol. 2009;39(8):877–82. Epub 2009/07/25. PubMed PMID: 19630138.

3. Jones JL, Dubey JP. Foodborne toxoplasmosis. Clin Infect Dis. 2012;55(6):845–51. Epub 20120522. doi: 10.1093/cid/cis508. PubMed PMID: 22618566.

4. Blader IJ, Coleman BI, Chen CT, Gubbels MJ. Lytic Cycle of *Toxoplasma gondii*: 15 Years Later. Annu Rev Microbiol. 2015;69:463–85. doi: 10.1146/annurev-micro-091014-104100. PubMed PMID: 26332089; PubMed Central PMCID: PMCPMC4659696.

5. Baum J, Papenfuss AT, Baum B, Speed TP, Cowman AF. Regulation of apicomplexan actin-based motility. Nat Rev Microbiol. 2006;4(8):621–8. doi: 10.1038/nrmicro1465. PubMed PMID: 16845432.

6. Lyons RE, McLeod R, Roberts CW. *Toxoplasma gondii* tachyzoite-bradyzoite interconversion. Trends Parasitol. 2002;18(5):198–201. PubMed PMID: 11983592.

7. Guo M, Mishra A, Buchanan RL, Dubey JP, Hill DE, Gamble HR, et al. A Systematic Meta-Analysis of *Toxoplasma gondii* Prevalence in Food Animals in the United States. Foodborne Pathog Dis. 2016;13(3):109–18. doi: 10.1089/fpd.2015.2070. PubMed PMID: 26854596.

8. Medlock MD, Tilleli JT, Pearl GS. Congenital cardiac toxoplasmosis in a newborn with acquired immunodeficiency syndrome. Pediatr Infect Dis J. 1990;9(2):129–32.

9. Lago EG, Conrado GS, Piccoli CS, Carvalho RL, Bender AL. Toxoplasma gondii antibody profile in HIV-infected pregnant women and the risk of congenital toxoplasmosis. Eur J Clin Microbiol Infect Dis. 2009;28(4):345–51. Epub 2008/10/16. doi: 10.1007/s10096-008-0631-2. PubMed PMID: 18855029.

10. Grant IH, Gold JW, Rosenblum M, Niedzwiecki D, Armstrong D. *Toxoplasma gondii* serology in HIV-infected patients: the development of central nervous system toxoplasmosis in AIDS. Aids. 1990;4(6):519–21.

11. Wang ZD, Wang SC, Liu HH, Ma HY, Li ZY, Wei F, et al. Prevalence and burden of Toxoplasma gondii infection in HIV-infected people: a systematic review and meta-analysis. Lancet HIV. 2017;4(4):e177–e88. Epub 2017/02/06. doi: 10.1016/S2352-3018(17)30005-X. PubMed PMID: 28159548.

12. van der Zypen E, Piekarski G. [Endodyogeny in Toxoplasma gondii. A morphological analysis]. Z Parasitenkd. 1967;29(1):15–35. Epub 1967/08/08. doi: 10.1007/BF00328836. PubMed PMID: 5601171.

13. Wohlfert EA, Blader IJ, Wilson EH. Brains and Brawn: Toxoplasma Infections of the Central Nervous System and Skeletal Muscle. Trends Parasitol. 2017;33(7):519–31. doi: 10.1016/j.pt.2017.04.001. PubMed PMID: 28483381; PubMed Central PMCID: PMCPMC5549945.

14. Dubey JP, Miller NL, Frenkel JK. The *Toxoplasma gondii* oocyst from cat feces. J Exp Med. 1970;132(4):636–62.

15. Ferguson DJ. Use of molecular and ultrastructural markers to evaluate stage conversion of Toxoplasma gondii in both the intermediate and definitive host. Int J Parasitol. 2004;34(3):347–60. doi: 10.1016/j.ijpara.2003.11.024. PubMed PMID: 15003495.

16. Speer CA, Clark S, Dubey JP. Ultrastructure of the oocysts, sporocysts, and sporozoites of *Toxoplasma gondii*. Journal of Parasitology. 1998;84(3):505–12.

17. Speer CA, Dubey JP. Ultrastructure of early stages of infections in mice fed *Toxoplasma gondii* oocysts. Parasitology. 1998;116(Pt 1)(6):35–42.

18. Zhu S, Shapiro K, VanWormer E. Dynamics and epidemiology of Toxoplasma gondii oocyst shedding in domestic and wild felids. Transbound Emerg Dis. 2022;69(5):2412–23. Epub 20210715. doi: 10.1111/tbed.14197. PubMed PMID: 34153160.

19. Vanwormer E, Conrad PA, Miller MA, Melli AC, Carpenter TE, Mazet JA. Toxoplasma gondii, source to sea: higher contribution of domestic felids to terrestrial parasite loading despite lower infection prevalence. Ecohealth. 2013;10(3):277–89. Epub 20130919. doi: 10.1007/s10393-013-0859-x. PubMed PMID: 24048652.

20. Hehl AB, Basso WU, Lippuner C, Ramakrishnan C, Okoniewski M, Walker RA, et al. Asexual expansion of Toxoplasma gondii merozoites is distinct from tachyzoites and entails expression of non-overlapping gene families to attach, invade, and replicate within feline enterocytes. BMC Genomics. 2015;16(1):66. Epub 20150213. doi: 10.1186/s12864-015-1225-x. PubMed PMID: 25757795; PubMed Central PMCID: PMCPMC4340605.

21. Farhat DC, Swale C, Dard C, Cannella D, Ortet P, Barakat M, et al. A MORC-driven transcriptional switch controls Toxoplasma developmental trajectories and sexual commitment. Nat Microbiol. 2020;5(4):570–83. Epub 20200224. doi: 10.1038/s41564-020-0674-4. PubMed PMID: 32094587; PubMed Central PMCID: PMCPMC7104380.

22. Augusto L, Wek RC, Sullivan WJ, Jr. Host sensing and signal transduction during Toxoplasma stage conversion. Mol Microbiol. 2021;115(5):839–48. Epub 20201121. doi: 10.1111/mmi.14634. PubMed PMID: 33118234; PubMed Central PMCID: PMCPMC9364677.

23. Martorelli Di Genova B, Wilson SK, Dubey JP, Knoll LJ. Intestinal delta-6-desaturase activity determines host range for Toxoplasma sexual reproduction. PLoS Biol. 2019;17(8):e3000364. Epub 20190820. doi: 10.1371/journal.pbio.3000364. PubMed PMID: 31430281; PubMed Central PMCID: PMCPMC6701743.

24. Bassermann F, Eichner R, Pagano M. The ubiquitin proteasome system - implications for cell cycle control and the targeted treatment of cancer. Biochim Biophys Acta. 2014;1843(1):150–62. Epub 20130301. doi: 10.1016/j.bbamcr.2013.02.028. PubMed PMID: 23466868; PubMed Central PMCID: PMCPMC3694769.

25. Silmon de Monerri NC, Yakubu RR, Chen AL, Bradley PJ, Nieves E, Weiss LM, et al. The Ubiquitin Proteome of *Toxoplasma gondii* Reveals Roles for Protein Ubiquitination in Cell-Cycle Transitions. Cell Host Microbe. 2015;18(5):621–33. doi: 10.1016/j.chom.2015.10.014. PubMed PMID: 26567513; PubMed Central PMCID: PMCPMC4968887.

26. Dogan T, Gnad F, Chan J, Phu L, Young A, Chen MJ, et al. Role of the E3 ubiquitin ligase RNF157 as a novel downstream effector linking PI3K and MAPK signaling pathways to the cell cycle. J Biol Chem. 2017;292(35):14311–24. Epub 2017/06/29. doi: 10.1074/jbc.M117.792754. PubMed PMID: 28655764; PubMed Central PMCID: PMCPMC5582827.

27. Liu Q, Tang Y, Chen L, Liu N, Lang F, Liu H, et al. E3 Ligase SCFbetaTrCP-induced DYRK1A Protein Degradation Is Essential for Cell Cycle Progression in HEK293 Cells. J Biol Chem. 2016;291(51):26399–409. Epub 2016/11/04. doi: 10.1074/jbc.M116.717553. PubMed PMID: 27807027; PubMed Central PMCID: PMCPMC5159501.

28. Johansson P, Jeffery J, Al-Ejeh F, Schulz RB, Callen DF, Kumar R, et al. SCF-FBXO31 E3 ligase targets DNA replication factor Cdt1 for proteolysis in the G2 phase of cell cycle to prevent re-replication. J Biol Chem. 2014;289(26):18514–25. Epub 2014/05/16. doi: 10.1074/jbc.M114.559930. PubMed PMID: 24828503; PubMed Central PMCID: PMCPMC4140253.

29. Galan JM, Wiederkehr A, Seol JH, Haguenauer-Tsapis R, Deshaies RJ, Riezman H, et al. Skp1p and the F-box protein Rcy1p form a non-SCF complex involved in recycling of the SNARE Snc1p in yeast. Mol Cell Biol. 2001;21(9):3105–17. doi: 10.1128/MCB.21.9.3105-3117.2001. PubMed PMID: 11287615; PubMed Central PMCID: PMCPMC86938.

30. Nelson DE, Randle SJ, Laman H. Beyond ubiquitination: the atypical functions of Fbxo7 and other F-box proteins. Open Biol. 2013;3(10):130131. doi: 10.1098/rsob.130131. PubMed PMID: 24107298; PubMed Central PMCID: PMCPMC3814724.

31. Wang Z, Liu P, Inuzuka H, Wei W. Roles of F-box proteins in cancer. Nat Rev Cancer. 2014;14(4):233–47. doi: 10.1038/nrc3700. PubMed PMID: 24658274; PubMed Central PMCID: PMCPMC4306233.

32. Lisztwan J, Marti A, Sutterluty H, Gstaiger M, Wirbelauer C, Krek W. Association of human CUL-1 and ubiquitin-conjugating enzyme CDC34 with the F-box protein p45(SKP2): evidence for evolutionary conservation in the subunit composition of the CDC34-SCF pathway. EMBO J. 1998;17(2):368–83. Epub 1998/02/28. doi: 10.1093/emboj/17.2.368. PubMed PMID: 9430629; PubMed Central PMCID: PMCPMC1170388.

33. Brunson LE, Dixon C, Kozubowski L, Mathias N. The amino-terminal portion of the F-box protein Met30p mediates its nuclear import and assimilation into an SCF complex. J Biol Chem. 2004;279(8):6674–82. Epub 2003/12/09. doi: 10.1074/jbc.M308875200. PubMed PMID: 14660673.

34. Gomi K, Sasaki A, Itoh H, Ueguchi-Tanaka M, Ashikari M, Kitano H, et al. GID2, an F-box subunit of the SCF E3 complex, specifically interacts with phosphorylated SLR1 protein and regulates the gibberellin-dependent degradation of SLR1 in rice. Plant J. 2004;37(4):626–34. Epub 2004/02/06. PubMed PMID: 14756772.

35. Sun L, Shi L, Li W, Yu W, Liang J, Zhang H, et al. JFK, a Kelch domain-containing F-box protein, links the SCF complex to p53 regulation. Proc Natl Acad Sci U S A. 2009;106(25):10195–200. Epub 2009/06/11. doi: 10.1073/pnas.0901864106. PubMed PMID: 19509332; PubMed Central PMCID: PMCPMC2700892.

36. Liu Y, Lear T, Zhao Y, Zhao J, Zou C, Chen BB, et al. F-box protein Fbxl18 mediates polyubiquitylation and proteasomal degradation of the pro-apoptotic SCF subunit Fbxl7. Cell Death Dis. 2015;6:e1630. Epub 2015/02/06. doi: 10.1038/cddis.2014.585. PubMed PMID: 25654763; PubMed Central PMCID: PMCPMC4669792.

37. Zheng N, Wang Z, Wei W. Ubiquitination-mediated degradation of cell cycle-related proteins by F-box proteins. Int J Biochem Cell Biol. 2016;73:99–110. doi: 10.1016/j.biocel.2016.02.005. PubMed PMID: 26860958; PubMed Central PMCID: PMCPMC4798898.

38. Mudhasani R, Tran JP, Retterer C, Kota KP, Whitehouse CA, Bavari S. Protein Kinase R Degradation Is Essential for Rift Valley Fever Virus Infection and Is Regulated by SKP1-CUL1-F-box (SCF)FBXW11-NSs E3 Ligase. PLoS Pathog. 2016;12(2):e1005437. Epub 2016/02/03. doi: 10.1371/journal.ppat.1005437. PubMed PMID: 26837067; PubMed Central PMCID: PMCPMC4737497.

39. Sidik SM, Huet D, Ganesan SM, Huynh MH, Wang T, Nasamu AS, et al. A Genome-wide CRISPR Screen in *Toxoplasma* Identifies Essential Apicomplexan Genes. Cell. 2016;166(6):1423–35 e12. doi: 10.1016/j.cell.2016.08.019. PubMed PMID: 27594426; PubMed Central PMCID: PMCPMC5017925.

40. Baptista CG, Lis A, Deng B, Gas-Pascual E, Dittmar A, Sigurdson W, et al. Toxoplasma F-box protein 1 is required for daughter cell scaffold function during parasite replication. PLoS Pathog. 2019;15(7):e1007946. doi: 10.1371/journal.ppat.1007946. PubMed PMID: 31348812; PubMed Central PMCID: PMCPMC6685633.

41. Sheiner L, Demerly JL, Poulsen N, Beatty WL, Lucas O, Behnke MS, et al. A systematic screen to discover and analyze apicoplast proteins identifies a conserved and essential protein import factor. PLoS Pathog. 2011;7(12):e1002392. Epub 2011/12/07. doi: 10.1371/journal.ppat.1002392. PubMed PMID: 22144892; PubMed Central PMCID: PMC3228799.

42. Sheffield HG, Melton ML. The fine structure and reproduction of *Toxoplasma gondii*. J Parasitol. 1968;54(2):209–26. PubMed PMID: 5647101.

43. Behnke MS, Wootton JC, Lehmann MM, Radke JB, Lucas O, Nawas J, et al. Coordinated progression through two subtranscriptomes underlies the tachyzoite cycle of *Toxoplasma gondii*. PLoS One. 2010;5(8):e12354. Epub 2010/09/25. doi: 10.1371/journal.pone.0012354. PubMed PMID: 20865045; PubMed Central PMCID: PMC2928733.

44. Suvorova ES, Francia M, Striepen B, White MW. A novel bipartite centrosome coordinates the apicomplexan cell cycle. PLoS Biol. 2015;13(3):e1002093. doi: 10.1371/journal.pbio.1002093. PubMed PMID: 25734885; PubMed Central PMCID: PMCPMC4348508.

45. Suvorova ES, Croken M, Kratzer S, Ting LM, Conde de Felipe M, Balu B, et al. Discovery of a splicing regulator required for cell cycle progression. PLoS Genet. 2013;9(2):e1003305. Epub 20130221. doi: 10.1371/journal.pgen.1003305. PubMed PMID: 23437009; PubMed Central PMCID: PMCPMC3578776.

46. Martins-Duarte ES, Carias M, Vommaro R, Surolia N, de Souza W. Apicoplast fatty acid synthesis is essential for pellicle formation at the end of cytokinesis in Toxoplasma gondii. J Cell Sci. 2016;129(17):3320–31. Epub 2016/07/28. doi: 10.1242/jcs.185223. PubMed PMID: 27457282.

47. Mazumdar J, E HW, Masek K, C AH, Striepen B. Apicoplast fatty acid synthesis is essential for organelle biogenesis and parasite survival in Toxoplasma gondii. Proc Natl Acad Sci U S A. 2006;103(35):13192–7. Epub 2006/08/22. doi: 10.1073/pnas.0603391103. PubMed PMID: 16920791; PubMed Central PMCID: PMCPMC1559775.

48. Striepen B, Crawford MJ, Shaw MK, Tilney LG, Seeber F, Roos DS. The plastid of *Toxoplasma gondii* is divided by association with the centrosomes. J Cell Biol. 2000;151(7):1423–34. Epub 2001/01/03. PubMed PMID: 11134072; PubMed Central PMCID: PMC2150670.

49. Amberg-Johnson K, Yeh E. Host Cell Metabolism Contributes to Delayed-Death Kinetics of Apicoplast Inhibitors in Toxoplasma gondii. Antimicrob Agents Chemother. 2019;63(2). Epub 2018/11/21. doi: 10.1128/AAC.01646-18. PubMed PMID: 30455243; PubMed Central PMCID: PMCPMC6355570.

50. Rolston KV. Treatment of acute toxoplasmosis with oral clindamycin. Eur J Clin Microbiol Infect Dis. 1991;10(3):181–3. Epub 1991/03/01. doi: 10.1007/BF01964456. PubMed PMID: 2060524.

51. Camps M, Arrizabalaga G, Boothroyd J. An rRNA mutation identifies the apicoplast as the target for clindamycin in *Toxoplasma gondii*. Mol Microbiol. 2002;43(5):1309–18. Epub 2002/03/29. PubMed PMID: 11918815.

52. Ramakrishnan C, Maier S, Walker RA, Rehrauer H, Joekel DE, Winiger RR, et al. An experimental genetically attenuated live vaccine to prevent transmission of Toxoplasma gondii by cats. Scientific reports. 2019;9(1):1474. Epub 20190206. doi: 10.1038/s41598-018-37671-8. PubMed PMID: 30728393; PubMed Central PMCID: PMCPMC6365665.

53. Behnke MS, Zhang TP, Dubey JP, Sibley LD. *Toxoplasma gondii* merozoite gene expression analysis with comparison to the life cycle discloses a unique expression state during enteric development. BMC Genomics. 2014;15:350. Epub 2014/06/03. doi: 10.1186/1471-2164-15-350. PubMed PMID: 24885521; PubMed Central PMCID: PMC4035076.

54. Fritz HM, Buchholz KR, Chen X, Durbin-Johnson B, Rocke DM, Conrad PA, et al. Transcriptomic analysis of toxoplasma development reveals many novel functions and structures specific to sporozoites and oocysts. PLoS One. 2012;7(2):e29998. Epub 20120213. doi: 10.1371/journal.pone.0029998. PubMed PMID: 22347997; PubMed Central PMCID: PMCPMC3278417.

55. Ramakrishnan C, Walker RA, Eichenberger RM, Hehl AB, Smith NC. The merozoite-specific protein, TgGRA11B, identified as a component of the Toxoplasma gondii parasitophorous vacuole in a tachyzoite expression model. Int J Parasitol. 2017;47(10-11):597–600. Epub 20170517. doi: 10.1016/j.ijpara.2017.04.001. PubMed PMID: 28526607.

56. Antunes AV, Shahinas M, Swale C, Farhat DC, Ramakrishnan C, Bruley C, et al. In vitro production of cat-restricted Toxoplasma pre-sexual stages by epigenetic reprogramming. bioRxiv. 2023. Epub 20230117. doi: 10.1101/2023.01.16.524187. PubMed PMID: 36711883; PubMed Central PMCID: PMCPMC9882236.

57. Saksouk N, Bhatti MM, Kieffer S, Smith AT, Musset K, Garin J, et al. Histone-modifying complexes regulate gene expression pertinent to the differentiation of the protozoan parasite *Toxoplasma gondii*. Mol Cell Biol. 2005;25(23):10301–14. Epub 2005/11/17. doi: 25/23/10301 [pii] 10.1128/MCB.25.23.10301-10314.2005. PubMed PMID: 16287846; PubMed Central PMCID: PMC1291236.

58. Bougdour A, Maubon D, Baldacci P, Ortet P, Bastien O, Bouillon A, et al. Drug inhibition of HDAC3 and epigenetic control of differentiation in Apicomplexa parasites. J Exp Med. 2009;206(4):953–66. Epub 2009/04/08. doi: jem.20082826 [pii] 10.1084/jem.20082826. PubMed PMID: 19349466; PubMed Central PMCID: PMC2715132.

59. Yan L, Lin M, Pan S, Assaraf YG, Wang ZW, Zhu X. Emerging roles of F-box proteins in cancer drug resistance. Drug Resist Updat. 2020;49:100673. Epub 20191217. doi: 10.1016/j.drup.2019.100673. PubMed PMID: 31877405.

60. Wang H, Maitra A, Wang H. The emerging roles of F-box proteins in pancreatic tumorigenesis. Semin Cancer Biol. 2016;36:88–94. Epub 20150916. doi: 10.1016/j.semcancer.2015.09.004. PubMed PMID: 26384530.

61. Gong J, Lv L, Huo J. Roles of F-box proteins in human digestive system tumors (Review). Int J Oncol. 2014;45(6):2199–207. Epub 20140929. doi: 10.3892/ijo.2014.2684. PubMed PMID: 25270675.

62. Yu H, Wu J, Xu N, Peng M. Roles of F-box proteins in plant hormone responses. Acta Biochim Biophys Sin (Shanghai). 2007;39(12):915–22. doi: 10.1111/j.1745-7270.2007.00358.x. PubMed PMID: 18064383.

63. Barth E, Hubler R, Baniahmad A, Marz M. The Evolution of COP9 Signalosome in Unicellular and Multicellular Organisms. Genome Biol Evol. 2016;8(4):1279–89. Epub 20160502. doi: 10.1093/gbe/evw073. PubMed PMID: 27044515; PubMed Central PMCID: PMCPMC4860701.

64. Schulze-Niemand E, Naumann M. The COP9 signalosome: A versatile regulatory hub of Cullin-RING ligases. Trends Biochem Sci. 2023;48(1):82–95. Epub 20220827. doi: 10.1016/j.tibs.2022.08.003. PubMed PMID: 36041947.

65. Ghosh S, Farr L, Singh A, Leaton LA, Padalia J, Shirley DA, et al. COP9 signalosome is an essential and druggable parasite target that regulates protein degradation. PLoS Pathog. 2020;16(9):e1008952. Epub 20200922. doi: 10.1371/journal.ppat.1008952. PubMed PMID: 32960936; PubMed Central PMCID: PMCPMC7531848.

66. Rosel D, Kimmel AR. The COP9 signalosome regulates cell proliferation of Dictyostelium discoideum. Eur J Cell Biol. 2006;85(9-10):1023–34. Epub 20060614. doi: 10.1016/j.ejcb.2006.04.006. PubMed PMID: 16781008.

67. Enchev RI, Scott DC, da Fonseca PC, Schreiber A, Monda JK, Schulman BA, et al. Structural basis for a reciprocal regulation between SCF and CSN. Cell Rep. 2012;2(3):616–27. Epub 20120906. doi: 10.1016/j.celrep.2012.08.019. PubMed PMID: 22959436; PubMed Central PMCID: PMCPMC3703508.

68. Cope GA, Suh GS, Aravind L, Schwarz SE, Zipursky SL, Koonin EV, et al. Role of predicted metalloprotease motif of Jab1/Csn5 in cleavage of Nedd8 from Cul1. Science. 2002;298(5593):608-11. Epub 20020815. doi: 10.1126/science.1075901. PubMed PMID: 12183637.

69. Jin D, Wu M, Li B, Bucker B, Keil P, Zhang S, et al. The COP9 Signalosome regulates seed germination by facilitating protein degradation of RGL2 and ABI5. PLoS Genet. 2018;14(2):e1007237. Epub 20180220. doi: 10.1371/journal.pgen.1007237. PubMed PMID: 29462139; PubMed Central PMCID: PMCPMC5834205.

70. Hartmann J, Hu K, He CY, Pelletier L, Roos DS, Warren G. Golgi and centrosome cycles in *Toxoplasma gondii*. Mol Biochem Parasitol. 2006;145(1):125–7. doi: 10.1016/j.molbiopara.2005.09.015. PubMed PMID: 16266757.

71. Nishi M, Hu K, Murray JM, Roos DS. Organellar dynamics during the cell cycle of *Toxoplasma gondii*. J Cell Sci. 2008;121(Pt 9):1559–68. Epub 2008/04/16. doi: 10.1242/jcs.021089. PubMed PMID: 18411248.

72. Skaar JR, D’Angiolella V, Pagan JK, Pagano M. SnapShot: F Box Proteins II. Cell. 2009;137(7):1358, e1. doi: 10.1016/j.cell.2009.05.040. PubMed PMID: 19563764.

73. Srivastava S, Holmes MJ, White MW, Sullivan WJ, Jr. Toxoplasma gondii AP2XII-2 Contributes to Transcriptional Repression for Sexual Commitment. mSphere. 2023;8(2):e0060622. Epub 20230214. doi: 10.1128/msphere.00606-22. PubMed PMID: 36786611; PubMed Central PMCID: PMCPMC10117075.

74. Wiley M, Sweeney KR, Chan DA, Brown KM, McMurtrey C, Howard EW, et al. *Toxoplasma gondii* activates hypoxia-inducible factor (HIF) by stabilizing the HIF-1alpha subunit via type I activin-like receptor kinase receptor signaling. J Biol Chem. 2010;285(35):26852–60. doi: 10.1074/jbc.M110.147041. PubMed PMID: 20581113; PubMed Central PMCID: PMCPMC2930684.

75. Alvarez CA, Suvorova ES. Checkpoints of apicomplexan cell division identified in *Toxoplasma gondii*. PLoS Pathog. 2017;13(7):e1006483. doi: 10.1371/journal.ppat.1006483. PubMed PMID: 28671988; PubMed Central PMCID: PMCPMC5510908.

76. Rahman K, Zhao P, Mandalasi M, van der Wel H, Wells L, Blader IJ, et al. The E3-ubiquitin ligase adaptor protein Skp1 is glycosylated by an evolutionarily conserved pathway that regulates protist growth and development. J Biol Chem. 2015;291(9):4268–80. Epub December, 30, 2015. doi: 10.1074/jbc.M115.703751. PubMed PMID: 26719340.

77. 1000GenomeProject. A map of human genome variation from population-scale sequencing. Nature. 2010;467(7319):1061–73.

78. Weber RJ, Li E, Bruty J, He S, Viant MR. MaConDa: a publicly accessible mass spectrometry contaminants database. Bioinformatics. 2012;28(21):2856–7. Epub 20120906. doi: 10.1093/bioinformatics/bts527. PubMed PMID: 22954629.

79. Pang Z, Zhou G, Ewald J, Chang L, Hacariz O, Basu N, et al. Using MetaboAnalyst 5.0 for LC-HRMS spectra processing, multi-omics integration and covariate adjustment of global metabolomics data. Nat Protoc. 2022;17(8):1735–61. Epub 20220617. doi: 10.1038/s41596-022-00710-w. PubMed PMID: 35715522.

80. Perez-Riverol Y, Csordas A, Bai J, Bernal-Llinares M, Hewapathirana S, Kundu DJ, et al. The PRIDE database and related tools and resources in 2019: improving support for quantification data. Nucleic Acids Res. 2019;47(D1):D442–D50. doi: 10.1093/nar/gky1106. PubMed PMID: 30395289; PubMed Central PMCID: PMCPMC6323896.

81. Ewels P, Magnusson M, Lundin S, Kaller M. MultiQC: summarize analysis results for multiple tools and samples in a single report. Bioinformatics. 2016;32(19):3047–8. Epub 20160616. doi: 10.1093/bioinformatics/btw354. PubMed PMID: 27312411; PubMed Central PMCID: PMCPMC5039924.

82. Kim D, Langmead B, Salzberg SL. HISAT: a fast spliced aligner with low memory requirements. Nat Methods. 2015;12(4):357–60. Epub 20150309. doi: 10.1038/nmeth.3317. PubMed PMID: 25751142; PubMed Central PMCID: PMCPMC4655817.

83. Love MI, Huber W, Anders S. Moderated estimation of fold change and dispersion for RNA-seq data with DESeq2. Genome Biol. 2014;15(12):550. doi: 10.1186/s13059-014-0550-8. PubMed PMID: 25516281; PubMed Central PMCID: PMCPMC4302049.

84. Singh P, Lonardi S, Liang Q, Vydyam P, Khabirova E, Fang T, et al. Babesia duncani multi-omics identifies virulence factors and drug targets. Nat Microbiol. 2023;8(5):845–59. Epub 20230413. doi: 10.1038/s41564-023-01360-8. PubMed PMID: 37055610; PubMed Central PMCID: PMCPMC10159843.

85. Bunnik EM, Venkat A, Shao J, McGovern KE, Batugedara G, Worth D, et al. Comparative 3D genome organization in apicomplexan parasites. Proc Natl Acad Sci U S A. 2019;116(8):3183–92. Epub 20190205. doi: 10.1073/pnas.1810815116. PubMed PMID: 30723152; PubMed Central PMCID: PMCPMC6386730.

86. Xia J, Venkat A, Bainbridge RE, Reese ML, Le Roch KG, Ay F, et al. Third-generation sequencing revises the molecular karyotype for Toxoplasma gondii and identifies emerging copy number variants in sexual recombinants. Genome Res. 2021;31(5):834–51. Epub 20210427. doi: 10.1101/gr.262816.120. PubMed PMID: 33906962; PubMed Central PMCID: PMCPMC8092015.

