## Supplementary material for "*Toxoplasma gondii* F-Box Protein L2 Silences Feline-Restricted Genes Necessary for Sexual Commitment": Fig S1

Figure S1. TGME49_313200, 10 exons, 832 aa

A. Predicted domain organization after HA-tagging. LRR = leucine-rich repeat, NLS = monopartite nuclear localization sequence. LRR3 and LRR4 are predicted by alphafold (see panel D), but are not identified by sequence prediction algorithms. Regions not labeled are poorly conserved.

B. Predicted amino acid sequence (from Toxo.db). Predicted domains are labeled and numbered as indicated. Underlined sequences were confirmed by mass spectrometry. Double underlined sequence represents the N-terminal 3X HA-tag.

C. Organization of amino acids 305-832 as predicted by alphaFold-2 (<https://alphafold.ebi.ac.uk/entry/S8EUA4>).

A.

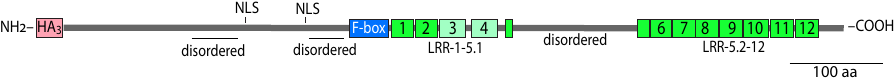

```
`
1  	MYPYDVPDYAGYPYDVPDYAYPYDVPDYAMLEARNASRADAARGPLRGTQGDEQTAERKE
61	TERARAQENADTFNTRSAKMSETVVNMREKEGREKGEQKEKKGQVAEEKKERRSERERPK

121	EKDLKSLFDSTDSDCLTSLETAKRALPKVEKKQMTILSFFSRPSSSSASSSPPSSSPPSS
													  NLS1
181	FSSSSSASSPSLLFRVPSALSRAASSTHTTEKRPTGPVDEKEEKKLLKRRRVCASEETDA
														  NLS2
241	LQGCARSFLAGMREAPRAGSASEANSKGEMSRDANVDSLHTPRLVEEHRKRRRTRLIQEH
														F-box
301	RAHNSSSSSSSSSSSSSSSSSSSSSSEKIELSPVGLSDLPEELLQQILDCCPKECLLVSH
		F-box				LRR1							LRR2
361	ALAAAVRRRRRVLRLTPLQCTEPRAEQFLSAIRSYPQLRLLEIVNAPWLSARFIENWANS
	   LRR2					  LRR3						    LRR4
421	KSSYFPQRLEVFVLRKCPRLTSRLARVLTTRLRRLRAIDLQDNSSLDYTSVVYLRTLPFL										LRR4			LRR5.1
481	ERVALGVSPSNTRGTSAHCNLTLCALLGPPSSSPTLLPSSTSSVLSSSSSSSLACSSASS

541	SVCPSTSTSPPSCLPSPNSTPVSSSSSSPRSPPSSLRWSGEKGEGREGEENEKRGSGSAL
													   LRR5.2	  LRR6
601	PRSKDEAGAGQAVREQRGETRAVDLLEKAQKQKQDQTAGSSSPPLKLLSLARCAALSSLA
			LRR6					LRR7
661	PLINVASTLEFLDLRGCCALDDSSAAVLASLANLQVLVLSDTGVTSTTVAAVAENCRRLE								LRR8			LRR9							LRR10
721	MLDVSRISRFSKEAALLLPLHLLRLTRLKLTKNSAVDDEVVRDCLSRLKRLALLDVSHCW										LRR11								LRR12
781	RVTSGFCVPPLPQNPDGMSLRRLGLFGCNVERQRVQDALRDAGAAKVQLSLHHELPMFDL

841	PCVYSNLPNLDLRCRTFIEGC*
```

D. Predicted organization of amino acids 305-832 (alphaFold-2). The F-box-like region is in red, and the 12 predicted LRRs are in cyan. Note that LRR5 is split by a long loop, not shown.

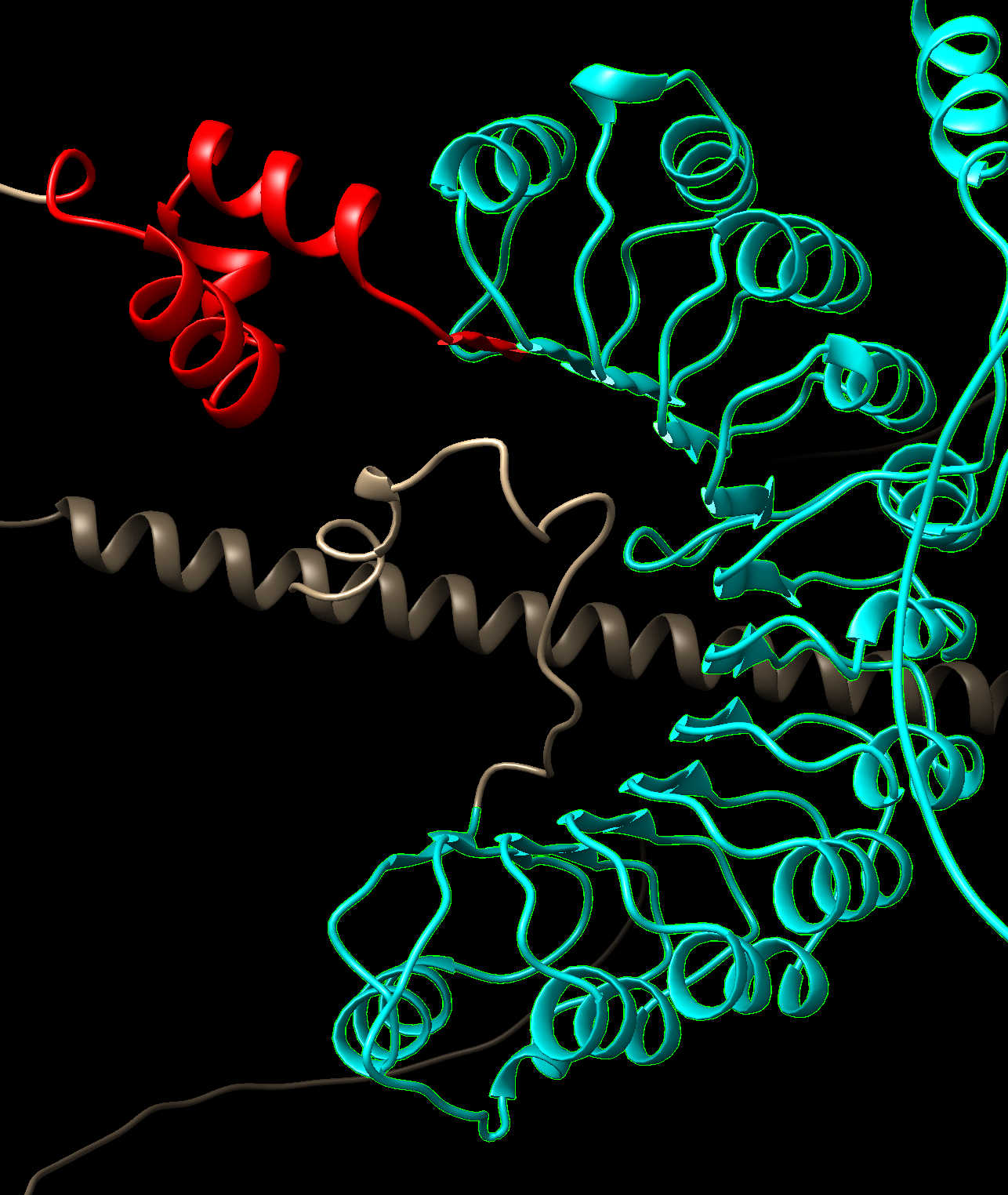
