## Supplementary figures and images for "*Toxoplasma gondii* F-Box Protein L2 Silences Feline-Restricted Genes Necessary for Sexual Commitment"

### Fig S2

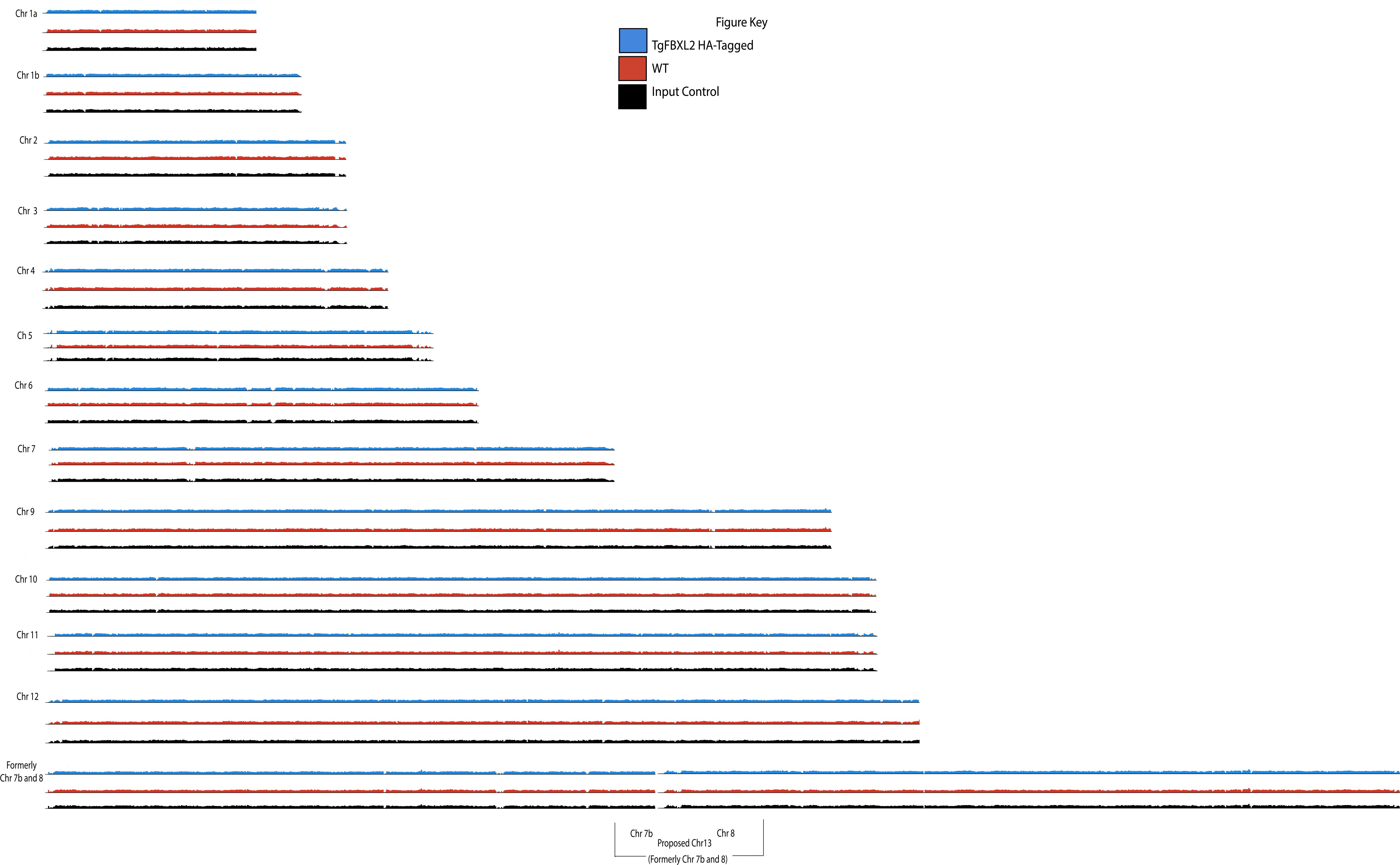
