## Supplementary material for "*Toxoplasma gondii* F-Box Protein L2 Silences Feline-Restricted Genes Necessary for Sexual Commitment": Fig S3

### Slide 1
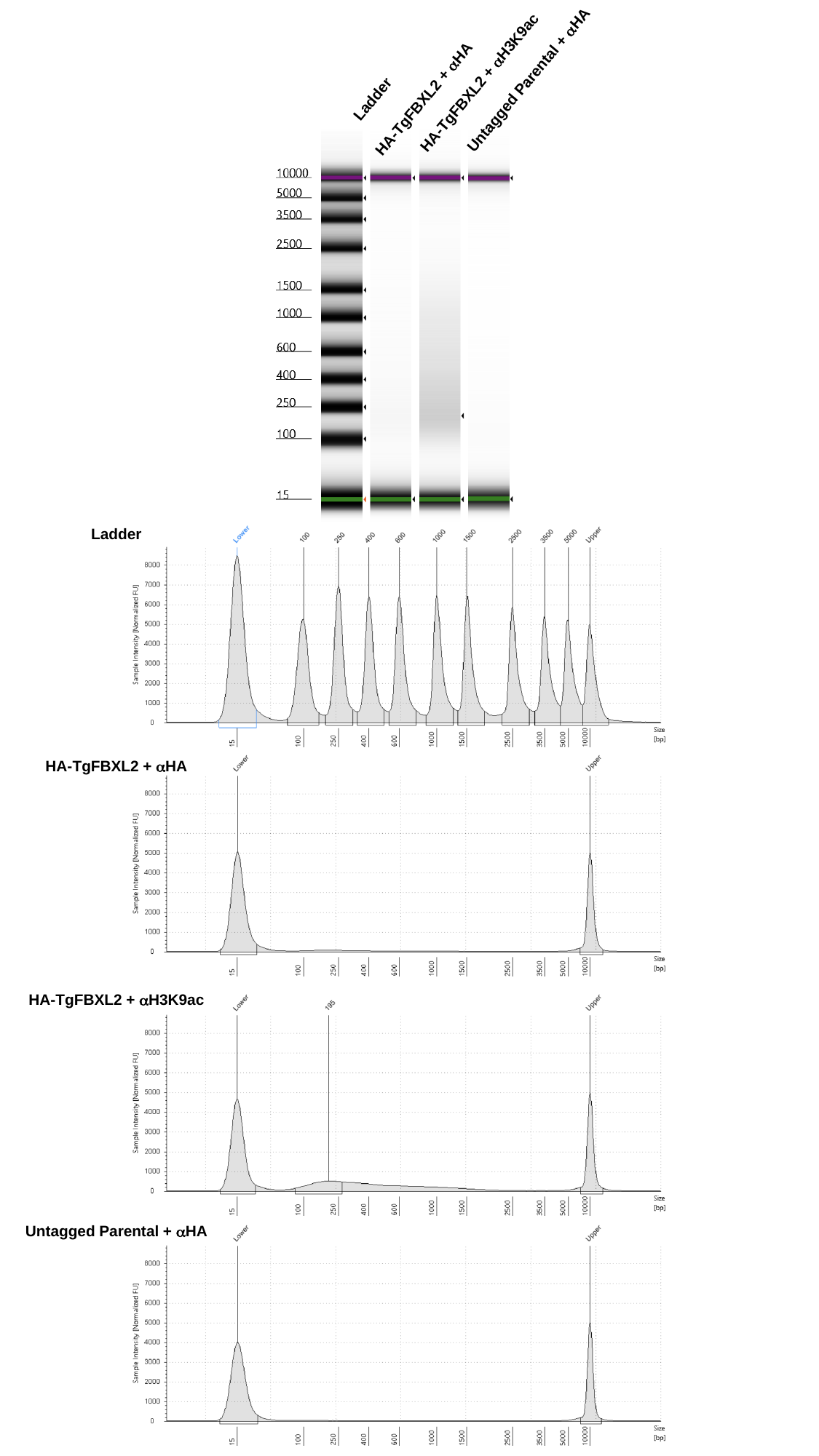

Untagged Parental + αHA
HA-TgFBXL2 + αH3K9ac
Ladder
HA-TgFBXL2 + αHA
Ladder
HA-TgFBXL2 + αHA
HA-TgFBXL2 + αH3K9ac
Untagged Parental + αHA
